# THE MULTIFUNCTIONAL PROTEIN E4F1 REGULATES MELANOCYTIC FUNCTIONS IN MICE THROUGH PROTEIN TRANSLATION AND MITF EXPRESSION

**DOI:** 10.64898/2026.10.05.756698

**Authors:** M. Rousseau, M. Houdeville, RL. Henry, PF. Roux, B. Seyran, L. Papon, J. Florido, C. Ferraro Peyret, M. Di Michele, B. Encislai, P. Dupré, N. Pirot, F. Bernex, A. Seguin, L. Rubio, A. Torro, A. Choquet, C. Chaveroux, L. Larue, A. David, L. Le Cam, M. Lacroix

## Abstract

The multifunctional protein E4 transcription factor 1 (E4F1) is an essential regulator of skin homeostasis, playing a critical role in epidermal stem cell maintenance and keratinocyte differentiation. However, the functions of E4F1 in other skin cell type remain poorly investigated. Here, we describe its role in melanogenesis, the physiological process for synthesis of melanin pigments. Using genetically engineered mouse models that enabled conditional inactivation of *E4f1* specifically in melanocytes, we demonstrated that *E4f1* loss induces severe pigmentation defects without affecting melanocyte survival nor differentiation. At the molecular level, *E4f1* depletion reduces the expression of its direct transcriptional target genes involved in protein translation, including *Elp3* and *Eef1g,* leading to translation alterations and consequently the induction of the Integrated Stress Response (ISR). This response resulted in microphthalmia-associated transcription factor (MITF) downregulation, inducing a decrease in the expression of enzymes involved in melanin synthesis. Consistent with these findings, *Elp3* or *Eef1g* depletion recapitulated the decrease in MITF levels in vitro, and pharmacological inhibition of ISR partially rescued melanin content in vivo, confirming a key role of the E4F1-protein translation-ISR cascade in melanogenesis. Together, our data identify E4F1-associated translation functions as a key regulator of the melanocytic lineage.

## INTRODUCTION

Melanogenesis is a physiological process responsible for synthesis of melanin pigments which occurs in melanocytes, a specialized cell type of the skin that transfer melanin to the surrounding keratinocytes to protect the skin from UV-induced DNA damages [1–3]. The amount, type and distribution of melanin determine hair, skin, and eye color [4,5]. Melanogenesis involves pigment-producing enzymes: tyrosinase (TYR), tyrosinase-related protein 1 (TYRP1) and dopachrome tautomerase (DCT), also known as tyrosinase-related protein 2 (TYRP2) [6]. The transcriptional expression of these enzymes is under the tight control of microphthalmia-associated transcription factor (MITF) [7–9]. *MITF* is a member of the Myc supergene family of basic helix-loop-helix leucine zipper (b-HLH-Zip) transcription factors and is known as the master regulator of the melanocytic lineage. It is essential for melanocyte development, function, survival and proliferation in both physiological and oncogenic context [10]. The first mutation of *Mitf* was discovered in the offspring of irradiated mice, which exhibited severe pigmentation loss due to a lack of neural crest-derived melanocytes, as well as microphtalmia and deafness [11]. Since, over 20 mutations in the *Mitf* gene have been identified in mice, all impacting melanocyte functions to varying degrees [12]. In humans, mutation in the *MITF* gene give rise to Waardenburg type 2 (WS2) and Tietz syndromes, which are characterized by pigmentation and deafness disorders [13–16]. MITF expression is regulated by several transcription factors, including SOX10, CREB, PAX3 and LEF1 [17] and many signaling pathways have been shown to regulate its expression [18–22]. A recent study demonstrated that translation reprogramming upon nutrient starvation leads to MITF repression at both the translational and transcriptional levels through the induction of the key component of the integrated stress response (ISR), ATF4 [23].

E4F1 was originally identified as a cellular target of the viral oncoprotein E1A during serotype V adenoviral infection [24]. E4F1 is a multifunctional, ubiquitously expressed protein that exhibits both transcriptional [24,25] and atypical ubiquitin E3 ligase activities [26]. It has been described as a key regulator of the p53 pathway: E4F1-mediated ubiquitylation of p53 modulates its transcriptional activities independently of its degradation [26]. In addition, E4F1 interacts directly with other components of the p53 pathway, including p14ARF [27] and BMI1 [28], and is a key regulator of the checkpoint protein CHK1 [29]. E4F1 was also identified as a central regulator of cellular metabolism, controlling pyruvate dehydrogenase activity [30] as well as lipid metabolism [31]. More recently, E4F1 has been implicated in the control of protein translation fidelity through the regulation of the *Elp3* gene, which encodes the catalytic subunit of the Elongator complex [32]. In vivo, E4F1 has been identified as a major actor in embryonic development and adult tissue homeostasis [31–34]. Especially, E4F1 plays an essential role in skin homeostasis and epidermal stem cell maintenance, involving both the BMI1-ARF-p53 pathway and pyruvate metabolism [34–36]. Beyond its roles in keratinocytes, the functions of E4F1 in other skin cell types remains poorly studied.

Here, we report that *E4f1* displays major functions in the melanocytic lineage. Using a new mouse model enabling specific inactivation of *E4f1* in melanocytes, as well as in vitro immortalized cell lines, we identified a pivotal role of E4F1 in melanogenesis and melanocyte homeostasis through the regulation of MITF. Impaired MITF activity is linked to the control of E4F1-direct target genes involved in protein translation, namely *Elp3* and *Eef1g.* Their downregulation leads to protein translation defects that induces an Integrated Stress Response (ISR) mediated by ATF4, a member of the ATF family which drives *Mitf* mRNA downregulation. Consistently, *Elp3* and *Eef1g* depletion recapitulated MITF decrease, and pharmacological inhibition of ISR partially rescued melanin content, confirming a key role of the protein translation in melanogenesis process.

## RESULTS

### Specific *E4f1* inactivation in melanocytes results in hair and skin pigmentation defects without impacting melanocytes survival nor differentiation

Using an *E4f1* whole-body conditional KO (cKO) mouse model (*E4f1*^lox^, RERT), we previously identified an essential role for E4F1 in adult skin homeostasis and epidermal stem cell maintenance [34–36]. Further characterization of this model revealed that topical application of tamoxifen on the tail of 8- to 12-weeks-old *E4f1^lox^*, RERT mice led to depigmentation of the tail skin (Fig. S1A-B). Immunohistochemistry (IHC) analyses of tail skin sections demonstrated a decreased expression of GP100, a marker of melanosomes, the organelles in which melanin synthesis takes place in melanocytes (Fig. S1C), suggesting a key role of E4F1 in melanocytes. To further investigate this role in the melanocytic lineage, we generated a new mouse model enabling *E4f1* inactivation specifically in melanocytes. We generated mice harboring one *E4f1* constitutive KO and one *E4f1* cKO allele, together with the TyrCreER^T2^ allele allowing expression of the tamoxifen (tam)-inducible CreER^T2^ fusion protein under the control of the melanocyte-specific tyrosinase promoter (*E4f1*^-/lox^, Tyr^CreERT2^ mice, and their corresponding *E4f1*^+/lox^, Tyr^CreERT2^ control animals, hereafter referred to as *E4f1*^(*Tyr*)KO^ and *E4f1*^(*Tyr*)CTL^ respectively) (Fig. S2A). A Cre-inducible ROSA26-Lox-Tomato-STOP-Lox-GFP (ROSA^mTmG^) reporter allele was also introduced in these compound mice to track recombined melanocytes (Fig. S2A). Topical applications of tamoxifen were performed on a shaved area of the back skin of 18 day-old *E4f1*^(*Tyr*)KO^ and *E4f1*^(*Tyr*)CTL^ mice to inactivate *E4f1* at the beginning of the first synchronous adult hair cycle (Fig. S2B). We validated the genetic inactivation of *E4f1* using in situ hybridization (RNAscope) and showed a strong decreased of *E4f1* mRNA in melanocytes of tam-treated *E4f1*^(*Tyr*)KO^ mice (Fig. 1A). 30 days after tam treatment, *E4f1*^(*Tyr*)KO^ mice showed severe defects in coat pigmentation and hair graying with absence of pigments in the hair shaft (Figs. 1B and S2C). These results were consistent with a strong decrease of melanin content in hairs in both *E4f1*^(*Tyr*)KO^ males and females (Figs. 1C and S2D). To determine whether this phenotype resulted from the loss of E4F1-deficient melanocytes, we quantified GFP-positive melanocytes in back skin sections prepared from *E4f1*^(*Tyr*)KO^ and *E4f1*^(*Tyr*)CTL^ mice. These analyses revealed no difference in the number of recombined melanocytes in the hair bulbs of *E4f1*^(*Tyr*)KO^ and *E4f1*^(*Tyr*)CTL^ (Fig. 1D) males and females (Fig. 1E). Consistent with the observed hair graying, melanin pigments detected in *E4f1*^(*Tyr*)CTL^ sections (in brown) was largely reduced in *E4f1*^(*Tyr*)KO^ skin samples, (Fig. 1D). Similarly, tam application on tail skin resulted in hypopigmentation in *E4f1*^(*Tyr*)KO^ mice (Fig. S2E), without impacting the number of interfollicular melanocytes in both males and females (Figs. S2F-G). To evaluate the differentiation status of the *E4f1* cKO melanocytes, the expression of the differentiation marker SOX10 was analyzed by immunofluorescence (IF) using back skin tissue sections and tail epidermal whole mounts. No difference in SOX10 expression was detected in melanocytes upon *E4f1* inactivation (Fig 1F-G).

**Figure 1:**
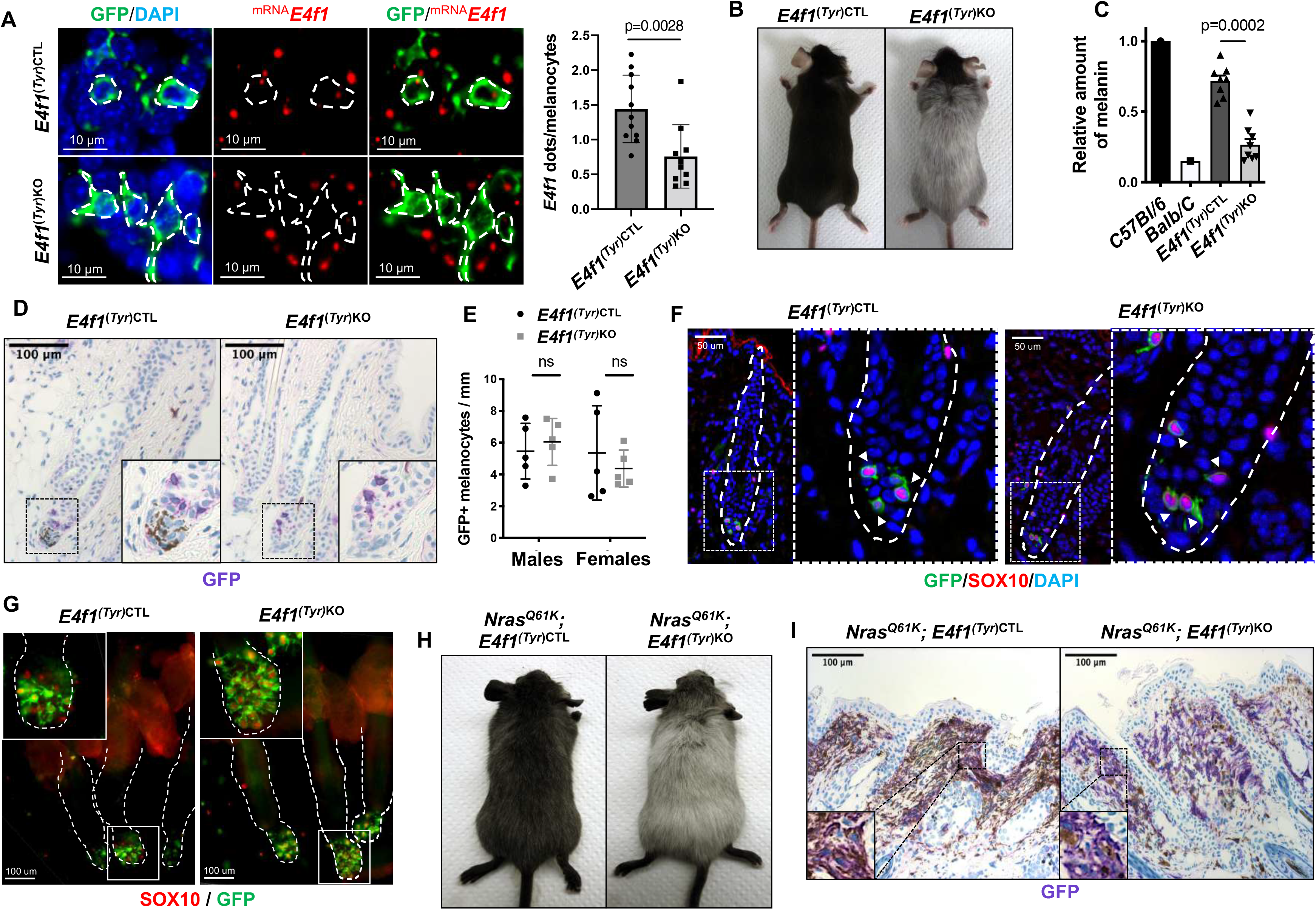
Specific *E4f1* inactivation in melanocytes results in hair and skin pigmentation defects. **(A)** Left panel: Microscopical images of E4f1 mRNA expression (red) in melanocytes (green) from *E4f1^(Tyr)CTL^*and *E4f1^(Tyr)KO^* back skin detected by RNAscope. Right panel: histobars showing the quantification of *E4f1* RNAscope signal (number of dots) per melanocytes. 60 to 100 GFP positive cells were counted and represented as the mean+/- SD of 11 individuals. Scale bar, 10 µm (**B)** Macroscopic observation of *E4f1^(Tyr)CTL^* et *E4f1^(Tyr)KO^* mice. *E4f1^(Tyr)KO^* animals exhibit hair graying compared to control littermates. **(C)** Melanin quantification in hairs from *E4f1^(Tyr^*^)*CTL*^ et *E4f1^(Tyr)KO^* (mean ± SEM, n=8). Hairs from C57Bl/6 and Balb/C mice were used as positive and negative control, respectively. **(D)** Immunohistochemistry analysis of the GFP reporter on back skin sections prepared from *E4f1^(Tyr)CTL^* et *E4f1^(Tyr)KO^* mice and showing follicular melanocytes in the hair bulb (arrow). Insets show images at higher magnification. Scale bar, 100 µm. **(E)** Quantification of GFP^+^ melanocytes per millimeter of back skin from males and females of *E4f1^(Tyr^*^)CTL^ et *E4f1^(Tyr^*^)*KO*^ mice (mean ± SEM, n=5 with 100 cells analyzed in each condition). **(F)** Immunofluorescence analysis of GFP (green) and SOX10 (red) on back skin section from *E4f1*^(*Tyr*)*CTL*^ et *E4f1*^(*Tyr*)*KO*^ mice. Sections were counterstained with DAPI (blue). Scale bar, 50 µm. **(G)** Microscopic three-dimensional image of whole mounts of tail epidermis prepared from *E4f1*^(*Tyr*)CTL^ et *E4f1*^(*Tyr*)KO^ mice. Sample were stained with GFP (green) and SOX 10 (red). Scale bar, 100µm. (**H**) Macroscopic observation of *Nras^Q61K^; E4f1*(*Tyr*)CTL et *Nras^Q61K^; E4f1*^(*Tyr*)KO^ mice. *Nras^Q61K^; E4f1*^(*Tyr*)KO^ animals exhibit hair graying compared to control littermates. **(I)** Immunohistochemistry analysis of the GFP reporter on back skin sections prepared from *Nras^Q61K^; E4f1*^(*Tyr*)CTL^ et *Nras^Q61K^; E4f1*^(*Tyr*)KO^ mice. Insets show images at higher magnification. Scale bar, 100 µm

We then wondered whether pigmentation defects due to *E4f1* inactivation could also occur in a hyperpigmented mouse model. To address this question, we crossed *E4f1*^lox^, TyrCreER^T2^ mice with strains allowing expression of the constitutively active mutated form of the *Nras^Q61K^* oncogene specifically in melanocytes (Tyr::*NRas*^Q61K^ model) (Fig. S2H). In this model, *E4f1* inactivation also led to strong coat pigmentation defects (Fig. 1H). IHC analysis of GFP-positive melanocytes on back skin sections showed an important decrease in melanin levels (in brown) in *Nras^Q61K^*; *E4f1*^(*Tyr*)KO^ even though melanocytes (in purple) were still detectable (Fig. 1I), indicating that *E4f1*inactivation in melanocytes prevents hyperpigmentation induced by the *Nras^Q61K^* oncogene without impacting melanocytes survival. Next, we addressed the potential role of the p53 tumor suppressor in the pigmentation defects observed E4F1-deficient melanocytes. p53 is a well-known actor of skin pigmentation in response to ultraviolet radiations [37] and previous studies established that E4F1 is a key regulator of the p53 pathway in keratinocytes [34]. We crossed *E4f1^lox^, TyrCreER^T2^* mice with *Trp53* KO strain to generate double compound KO animals (hereafter referred to as *E4f1*^(*Tyr*)CTL^; *p53*^KO^ and *E4f1*^(*Tyr*)KO^; *p53*^KO^). Tam-induced *E4f1* inactivation in these double KO mice resulted in coat pigmentation defects (Fig. S3A) and decreased melanin content (Fig. S3B) similar to those observed in *E4f1*^(*Tyr*)KO^ mice, indicating that the pigmentation defects caused by *E4f1* inactivation are not p53-dependent.

Taken together, these data show that *E4f1* inactivation does not affect melanocytes survival nor differentiation, and suggested that E4F1-associated pigmentation defects could result from a cell-autonomous impairment of the melanogenesis process.

### E4F1 regulates the expression of MITF and melanin synthesis enzymes

We next addressed the molecular mechanism by which *E4f1* regulates melanogenesis. For this purpose, murine pigmented and immortalized melanocytes, Melan-a cells, were transduced with lentiviral particles encoding a control shRNA (shCTL) or shRNA targeting *E4f1* mRNA (sh*E4f1*). First, we validated E4F1 protein depletion by immunoblotting in sh*E4f1* Melan-a cells (Fig 2A) and then determined their transcriptome by RNA sequencing (RNA-seq). Upon *E4f1* depletion, a total of 2725 genes were differentially expressed with a false discovery rate (FDR) inferior to 0.05, among which 1468 were down-regulated and 1257 were up-regulated (Figs. 2B-C and Supplemental Table S1). Among the most downregulated transcripts, we found *Neurl4*, *Fktn*, *Dlat*, *Bag5*, *Ipo8*, *Tube1*, *Pbrm1*, *Wdr7*, *Eef1g*, *Dnajc19* and *Taz*, which were previously identified as E4F1-direct target genes in mouse embryonic fibroblasts (MEFs) and embryonic stem (ES) cells [29]. In addition, gene set enrichment analysis (GSEA) indicated that the E4F1 signature was significantly repressed in sh*E4f1* Melan-a cells (Fig. 2D), and showing that the core transcriptional program controlled by *E4f1* in fibroblasts and neural cells is also regulated in melanocytes.

**Figure 2:**
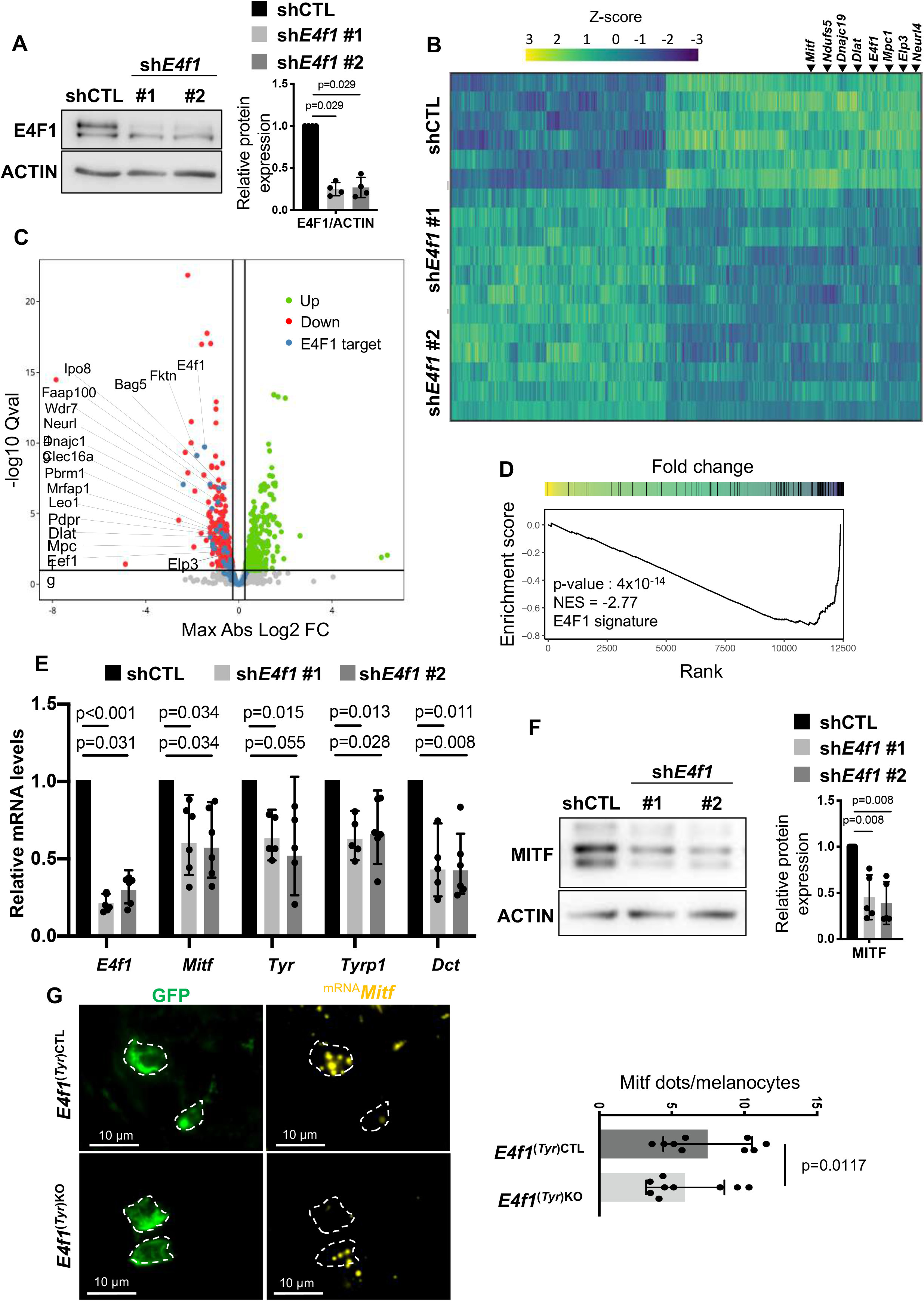
E4f1 regulates MITF and melanin synthesis enzymes expression. **(A)** Immunoblot analysis of E4F1 and ACTIN (loading control) protein levels in Melan-a cells, 10 days after transduction with lentiviral particles encoding an irrelevant shRNA (shCTL) or shRNA targeting *E4f1* mRNA (sh*E4f1*). **(B)** Heatmap depicting expression levels across conditions for genes differentially expressed upon E4f1 depletion (FDR corrected p-value >.1 and leading log2 fold-change > log2(1.5)). Data are expressed as row Z-scores collected from five biologically independent experiments per condition. **(C)** Volcano plots depicting the −log10 p-value as a function of the log2 fold-change in gene expression defined by a differential analysis conducted with EdgeRto highlight the effect of E4f1 depletion. Green and red dots highlight up and down regulated genes, respectively. Blue dots highlight genes annotated as E4F1 targets. **(D)** Evolution of score (ES) along ranking as defined with a Gene Set Enrichment Analysis (GSEA) comparing gene expression fold-change between *E4f1*-depleted and control cells and known E4F1 target genes. 1,000 permutations were performed and the associated Benjamini-Hochberg adjusted p-value and NES (Normalized Enrichment Score) is provided. **(E)** RT-qPCR analysis of *Mitf* and melanin synthesis enzymes mRNA levels in shCTL and sh*E4f1* Melan-a cells (mean +/- SEM, n=6). **(F)** Immunoblot analysis of MITF and ACTIN (loading control) protein levels in Melan-a cells, 10 days after transduction with lentiviral particles encoding an irrelevant ShRNA (shCTL) or shRNA targeting *E4f1* mRNA (sh*E4f1*). **(G)** Left panel: Microscopical images of *Mitf* mRNA expression (yellow) in melanocytes (green) from *E4f1^(Tyr)CTL^*and *E4f1^(Tyr)KO^* back skin detected by RNAscope. Scale bar, 10 µm. Right panel: Histobar showing the quantification of *Mitf* RNAscope signal (number of dots) per melanocytes.

Looking for candidate genes related to pigmentation, we identified the master regulator of melanogenesis, *Mitf*, among the significantly down-regulated transcripts in *E4F1*-depleted Melan-a cells. We confirmed the downregulation of MITF by immunoblotting and RT-qPCR analyses in sh*E4f1* Melan-A cells. Moreover, we also observed decreased expression of Tyrosinase (*Tyr*), *Tyrp1* and *Dct*, which are directly regulated by MITF (Fig. 2E-F). We next evaluated the relevance of this finding in vivo and analyzed the expression of *Mitf* mRNA using in situ hybridization on skin sections from *E4f1*^(*Tyr*)CTL^ and *E4f1*^(*Tyr*)KO^ mice. We showed that *Mitf* transcript levels were decreased in GFP-positive *E4f1*^(*Tyr*)KO^ melanocytes compared to control mice, confirming our in vitro data and suggesting that the pigmentation defect was due, at least in part, to the down-regulation of MITF (Fig. 2G). Taken together, these data demonstrated a key role for E4F1 in the transcriptional regulation of *Mitf* in melanocytes.

### *E4f1* regulates genes involved in protein translation that controls MITF expression in the melanocytic lineage

We then further characterized the molecular mechanism regulating MITF expression in E4F1-deficient melanocytes. Several signaling pathways have been shown to regulate MITF expression, and recent data demonstrated that translation reprogramming leads to MITF repression through the induction of the integrated stress response (ISR) [38]. Previous ChIP-seq experiments performed in murine embryonic fibroblasts and ES cells identified several genes involved in protein translation as potential direct target genes of E4F1 (Fig. 3A) [41]. Among these genes, we recently identified *Elp3*, encoding the catalytic subunit of the Elongator complex implicated in translation fidelity [32]; *Eef1g, Eif3d and Eif4b*, which encode proteins involved in translation elongation and initiation respectively. Interestingly, GSEA analysis highlighted that translation signatures (including Translation initiation and Translation elongation) were downregulated in sh*E4f1* Melan-a cells (Fig. 3B). Next, we analyzed the expression of these E4F1-direct target genes in *E4f1*-depleted Melan-a cells and showed a significant decrease of their mRNA and protein levels (Figs. 3C-D). We confirmed these results in vivo using in situ hybridization on skin sections prepared from *E4f1*^(*Tyr*)CTL^ and *E4f1*^(*Tyr*)KO^ mice. We showed that *Elp3 and Eef1g* transcripts were decreased in *E4f1*^(*Tyr*)KO^ melanocytes compared to control mice (Fig 3E).

**Figure 3:**
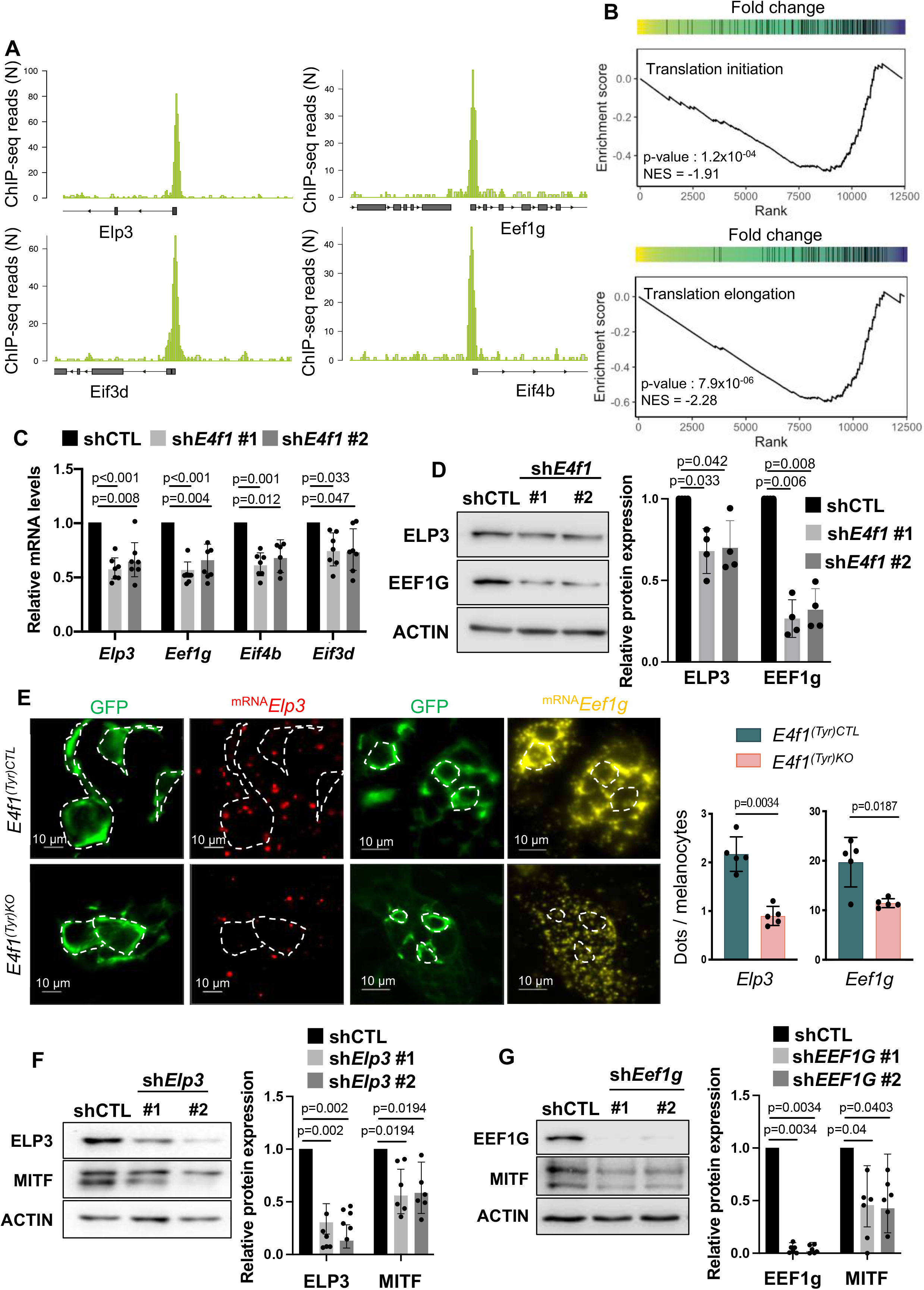
E4F1 regulates gene involved in translation in melanocyte. **(A)** E4F1 ChIP-seq read densities in transformed MEFs (tMEFs) at the *Elp3, Eef1g, Eif3d, Eif4b* locus. **(B)** Evolution of score (ES) along ranking as defined with a Gene Set Enrichment Analysis (GSEA) comparing gene expression fold-change involved in Translation initiation and Translation elongation between E4f1-depleted and control cells. 1,000 permutations were performed and the associated Benjamini-Hochberg adjusted p-value and NES (Normalized Enrichment Score) is provided. (**C**) RT-qPCR analysis of *Elp3, Eef1g, Eif4b and Eif3d* mRNA levels in shCTL and sh*E4f1* Melan-a cells (mean +/- SEM, n=7). (**D**) Left panel: Immunoblot analysis of ELP3, EEF1G and ACTIN (loading control) protein levels in shCTL and sh*E4f1* Melan-a cells, 10 days after transduction. Right panel: histobars represent the quantification of dots per GFP positive melanocytes performed on 5 independent experiments. (**E**) Left panel: Microscopical images of *Elp3* (red) and *Eef1g* mRNA expression (yellow) in melanocytes (green) from *E4f1^(Tyr)CTL^*and *E4f1^(Tyr)KO^* back skin detected by RNAscope. Scale bar, 10 µm. Right panel: histobars showing the quantification of *Elp3 and Eef1g* RNAscope signal (number of dots) per melanocytes. (**F**) Immunoblot analysis of ELP3, MITF and ACTIN (loading control) protein levels in Melan-a cells, 7 days after transduction with lentiviral particles encoding an irrelevant ShRNA (shCTL) or ShRNA targeting *Elp3* (sh*Elp3*). (**G**) Immunoblot analysis of EEF1G, MITF and ACTIN (loading control) protein levels in Melan-a cells, 7 days after transduction with lentiviral particles encoding an irrelevant ShRNA (shCTL) or shRNA targeting *Eef1g* (sh*Eef1g*).

To further strengthen the role of ELP3 and EEF1g in the E4F1-associated regulation of MITF expression, we evaluated the molecular consequences of their depletion in Melan-a cells. We demonstrated that shRNA-mediated knock-down of *Elp3* led to an impact on protein translation in puromycin assay together with a decreased expression of MITF at the protein levels (Figs. 3F and S4A). Interestingly, shRNA-mediated knock-down of *Eef1g* also recapitulated the decrease in protein translation and the downregulation of MITF protein levels (Figs. 3G and S4B).

Altogether, these data support the notion that *E4f1* inactivation leads to perturbed activity of the Elongator complex and of the translation initiation/elongation complexes that correlate with MITF down regulation.

### E4F1 depletion impacts on protein translation and induces an Integrated Stress Response (ISR)

To evaluate the consequences of E4F1-depletion on translation, we first performed a puromycin-incorporation assay in sh*E4f1* and shCTR Melan-A cells. This assay confirmed that E4F1 depletion has a strong impact on protein translation in melanocytes (Fig. 4A). To identify which mRNAs were impacted, we next performed polysome profiling experiments. We isolated total cytosolic mRNAs, as well as mRNAs from pooled light (1 to 3 ribosomes) and heavy (more than 3 ribosomes) polysomal fractions prepared from *E4f1*-depleted and control Melan-a cells at day 10, and then determined the abundance of these transcripts using next generation sequencing (Figs. 4B and S4C). We first focused on a group of genes exhibiting similar total transcript levels but differential mRNA abundance in either the light or the heavy fractions purified from *E4f1*-depleted Melan-a cells. Consistent with the potential role of ELP3 in E4F1-dependent translation defects, bioinformatic analysis of codons content showed that these differentially translated mRNAs are significantly enriched in two U_34_-codons (AAA, GAA) in comparison to transcripts of the whole transcriptome (Fig. 3C).

**Figure 4:**
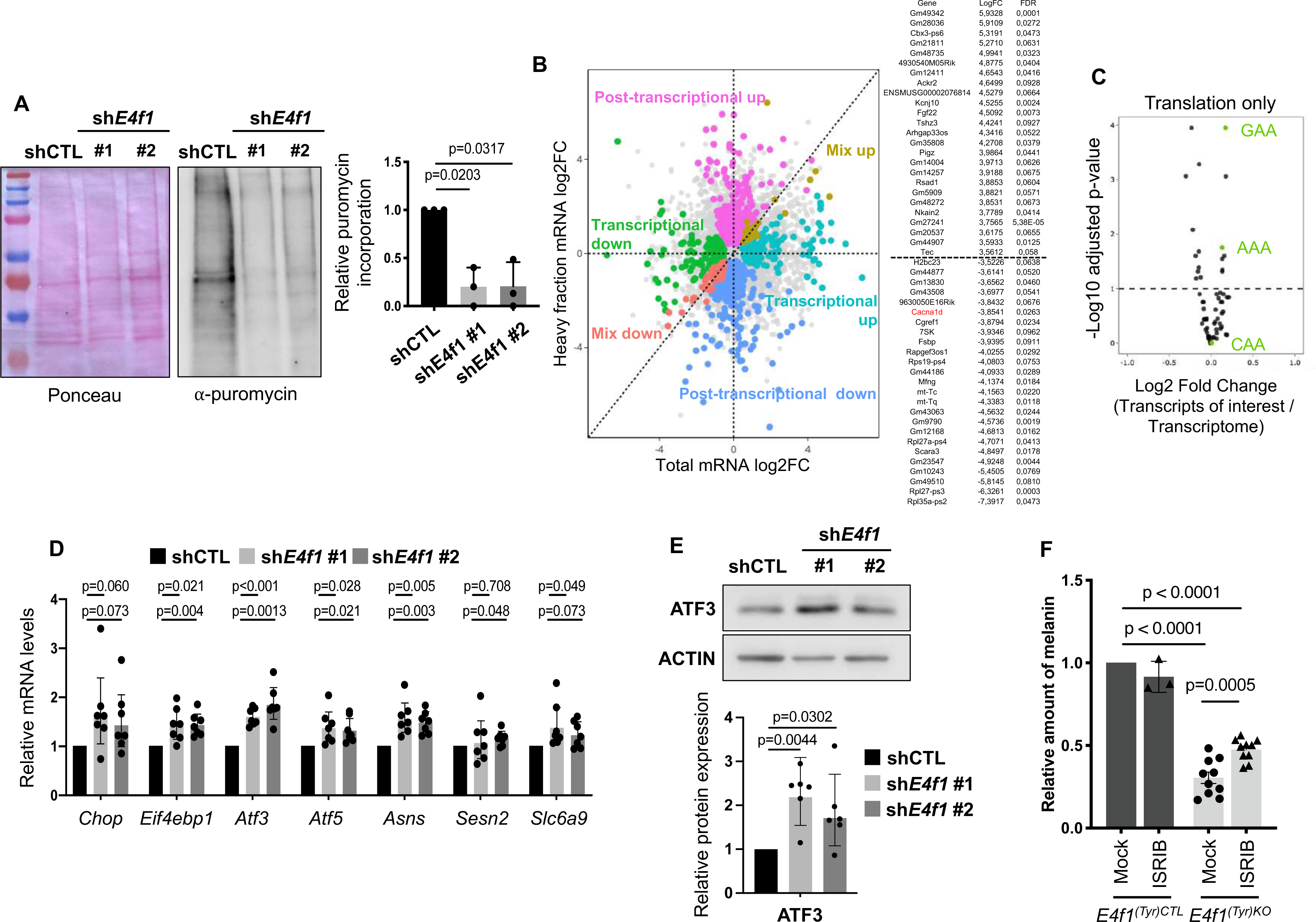
E4F1 depletion leads to translation defects and to an ISR in vitro and in vivo. **(A)** Puromycin assay in shCTL and sh*E4f1* Melan-a cells. Left panel: Immunoblot analysis of puromycin in Melan-a cells, 10 days after transduction with lentiviral particles encoding an irrelevant shRNA (shCTL) or shRNA targeting *E4f1* (sh*E4f1*). Right panel: histobars represent the quantification of immunoblots performed on 3 independent experiments. (**B**) Polysome profiling analysis of brains of shCTL and sh*E4f1* Melan-a cells (n=5). Dot plots represent transcript levels (Log_2_ fold change) between sh*E4f1* and shCTL Melan-a cells in the pooled heavy (more than 3 ribosomes, right panel) polysomal fractions, according to their abundance in the total cytosolic fraction. The lists indicate the 16 genes which transcript abundance is the most deregulated (log_2_FC) in the heavy polysomal fractions prepared from sh*E4f1* Melan-a cells but showing similar total transcript levels. (**C**) Codon enrichment in transcripts regulated at the post-transcriptional level in the brain of shCTL and sh*E4f1* Melan-a cells. U_34_-linked codons are indicated in green. Codon enrichment was assessed by χ^2^ tests. **(D)** Relative mRNA levels of genes involved in ER stress/UPR in shCTL and sh*E4f1* Melan-a cells (mean ± SEM, n=7). (**E**) Upper panel: Immunoblot analysis of ATF3 and ACTIN (loading control) in Melan-a cells 10 days after transduction with lentiviral particles encoding an irrelevant shRNA (shCTL) or shRNA targeting *E4f1* (sh*E4f1*). Lower panel: histobars represent the quantification of immunoblots performed on 6 independent experiments. (**F**) Relative amount of melanin in hair of *E4f1^(Tyr)CTL^*and *E4f1^(Tyr)KO^* back skin upon ISRIB treatment.

Furthermore, analysis of our RNAseq datasets pointed at the UPR response among the more significantly enriched pathways identified in E4F1-depleted Melan-A cells (Fig. S5). More precisely, we identified *Atf3*, *Atf5*, *Slc6a9*, *Asns* and *Eif4ebp1* transcripts as upregulated in *E4f1*-depleted Melan-a cells, and we confirmed the induction of these ATF-related genes by RT-qPCR (Fig. 4D), as well as the increase of ATF3 protein levels by immunoblotting (Fig. 4E). To confirm the importance of the ISR in the hair graying phenotype observed in *E4f1*^(*Tyr*)KO^ mice, we treated daily these mice with ISRIB, a selective ISR inhibitor, from day 18 to 51, and quantified the melanin content in the hair. ISRIB-treatment partially rescued the amount of melanin in *E4f1*^(*Tyr*)KO^ mice treated (Fig. 4F). Altogether, these data support a model where deregulation of E4F1-mediated control of genes involved in protein translation in melanocytes leads to the induction of an ISR, which in turn induces depigmentation.

## DISCUSSION

We previously identified E4F1 as essential regulator of skin homeostasis. *E4f1* inactivation leads to severe skin alterations due to impaired epidermal stem cell maintenance. However, a deeper characterization of *E4f1* whole body KO model revealed a potential role of E4F1 in melanocytes. Here, we describe a role of E4F1 in melanocyte homeostasis through the regulation of MITF. We show that E4F1 functions in melanocytes occurs through the control of protein translation since perturbation of *E4f1* expression leads to strong translation defects and the induction of an ISR response that alter MITF expression and melanocyte functions. Using genetically engineered mice lacking *E4f1* specifically in melanocytes, we show that *E4f1* inactivation induces severe coat and skin pigmentation defects. Although E4F1 has been shown to be essential for the survival of neural progenitors [32], immortalized and transformed cells [29, 39, 40], our data indicate that deletion of *E4f1* does not induce cell death in melanocytes. We confirmed this notion in a mouse model of hyper-pigmentation driven by mutated NRAS. Moreover, since we found that the expression of the differentiation marker SOX10 was not altered in *E4f1*-deficient mice, our data suggest that E4F1 plays a major intrinsic role in melanocyte functions and melanogenesis rather than impacting on cell survival or differentiation. Furthermore, despite the previously described role of the p53 tumor suppressor in UV-induced melanogenesis and its link with the E4F1 protein [37], we demonstrate that p53 inactivation does not rescue the pigmentation defects of *E4f1*^(*Tyr*)KO^ mice, suggesting that E4F1 functions in keratinocytes and melanocytes involve different molecular mechanisms.

Using immortalized melanocytes, we demonstrate that the E4F1-mediated pigmentation defects are cell autonomous and involve MITF and its target genes involved in melanogenesis (*Tyr*, *Tyrp1* and *Dct*). Several studies have shown that MITF can be directly repressed by the transcription factor ATF4, a key component of the ER stress and ISR responses [41] consequently to translation reprogramming [23, 38]. Interestingly, we recently identified E4F1 as a direct transcriptional regulator of *Elp3*, encoding the catalytic subunit of the Elongator complex [32]. This atypical acetyl-transferase acetylates a subset of tRNAs at the wobble position U_34_ to promote a stepwise cascade of tRNAs modifications that ensure translation fidelity at specific codons [42]. *E4f1* inactivation during neuronal development induced alteration of the activity of the Elongator complex, translation defects and activation of an ISR response mediated by the PERK-eiF2*a*-ATF4 pathway [32]. Similarly, we demonstrated that *E4f1*-depleted melanocytes also displayed a disruption of the Elongator complex but also of several other components of the translation machinery such as EEF1g and EIF4b leading to the induction of an ISR response that correlates with the downregulation of *Mitf.* Interestingly, we were able to recapitulate MITF downregulation associated to *E4f1* knock down by depleting either *Elp3* or *Eef1g* illustrating that E4F1 regulation of MITF and melanogenesis is mediated through several mechanisms impacting on protein translation. Although we identified a key role for E4F1 in this cascade in homeostatic conditions, it will be interesting to evaluate whether it is also involved in other situations such as nutritional stress.

Interestingly, ISR and UPR have been frequently associated with pigmentary disorders. Mutations responsible for albinism, in which melanocytes are unable to produce melanin, induced misfolding and accumulation in ER of the tyrosinase protein, leading to ER stress and UPR [43,44]. Unsolved UPR usually induces apoptosis [45]. However, albino melanocytes have been described as particularly resistant to UPR-mediated cell death [46,47], which would explain their survival upon *E4f1* inactivation in our model. Similarly, dilated endoplasmic reticulum, which are commonly associated with accumulation of misfolded proteins, has been observed in melanocytes of patients with vitiligo, a complex autoimmune disease characterized by absent or dysfunctional melanocytes [48]. Despite this notion remains to be confirmed, these data suggest that translation defects might be a common feature of pigmentary disorders. In addition, membrane lipid defects and in particular an altered distribution of cardiolipins, leading to the production of ROS, have been described in melanocyte mitochondria of patients with vitiligo [49]. Interestingly, we previously identified E4f1 as a transcriptional regulator of *Taz*, *Crls1* and *Dnajc19*, three genes directly involved in cardiolipin synthesis [29]. Although no *E4F1* mutation has been associated so far with pigmentary diseases in humans, studying the role of E4F1 in these pathologies could bring novel insights into the understanding of the mechanisms inducing melanocyte dysfunction.

Finally, beyond its physiological relevance, our finding that *E4f1* inactivation prevents NRAS-driven hyperpigmentation raises the possibility that this translation-ISR-MITF axis may also be relevant in the context of melanoma, where MITF plays a central and context-dependent role in tumor phenotype switching and therapy resistance. Given the established involvement of translational control and the ISR in cancer cell adaptation, our results suggest that the E4F1-ELP3/EEF1g-ISR-MITF axis identified here could represent a broader regulatory node worth investigating in melanoma biology.

## METHODS

### Generation of mouse models, Animal housing, Experimental treatment and genotyping

Generation of *E4f1* cKO mice was previously described [24]. These mice were first intercrossed with Cre-ER^T2^ knock-in mice (*RERT, Polr2a^tm1(cre/ERT2)Bbd^/J*) to generate *E4f1*^+/flox^, *RERT^KI/KI^*; *E4f1*^-/flox^, *RERT^KI/KI^* experimental groups (referred as to *E4f1*(*RERT*)CTL and *E4f1*(*RERT*)KO respectively). *E4f1* cKO mice were also intercrossed with TyrCreER^T2^ mice and mice carrying Rosa26^mTmG^ reporter (B6.129(Cg)-Gt(ROSA)26Sor^tm4(ACTB-tdTomato,-EGFP)Luo^/J) to generate *E4f1*^+/flox^, TyrCreER^T2^, mTmG^KI/+^ and *E4f1*^-/flox^, TyrCreER^T2^, mTmG^KI/+^ experimental groups (referred as to *E4f1*(*Tyr*)CTL and *E4f1*(*Tyr*)KO respectively). These mice were then intercrossed with p53KO strain (B6.129S2-*Trp53^tm1Tyj^*/J) to generate double KO E4f1/p53 compound mice (referred as to *E4f1*(*Tyr*)CTL; p53KO and *E4f1*(*Tyr*)KO; p53KO). *E4f1* cKO mice were also intercrossed either with mice allowing the constitutive expression of *TyrNras^Q61K^* (MGI:3768645).

Mice, generated in a mix 129Sv/J; C57Bl/6 background, were housed in a pathogen free barrier facility (temperature 22°C; 55% humidity; 12h cycle light/dark) in accordance with the ethic committee (Comité d’éthique pour l’Expérimentation Animale CEEA) for animal warfare (agreement number #17079-2018100514023945 v2). Mice were maintained under chow diet (A03, Safe) containing 22kcal% protein, 65kcal% carbohydrate and 13kcal% fat. Cre-mediated recombination of the *E4f1^flox^* allele was induced by topical application of tamoxifen (Sigma; 2mg/day/mouse diluted in ethanol for 5 consecutive days) on shaved back or tail skin of 8- to 12-weeks-old or 18-days-old animals for RERT and TyrCreER^T2^ models respectively.

TyrCreER^T2^ mice were treated with ISRIB (2.5 mg/kg/day, Sigma, SML0843) by intraperitoneal injections following tamoxifen application, 5 days a week.

Mice were genotyped by PCR on tail genomic DNA using GoTaq® G2 Flexi DNA Polymerase kit (Promega). Primers used for genotyping are listed in Supplemental Table S2.

### Melanin Extraction and Quantification

Melanin was extracted from dorsal hairs (1.5 mg) by alkalin treatment (1.5 ml of 1M NaOH, 4 hr at 85°C). Relative melanin content was determined by spectrophotometric measurement at 475 nm. Each hair sample was measured in triplicate and normalized with values obtained for C57Bl/6 hairs and compared to values obtained for Balb/C hairs.

### Immunohistochemistry, Immunofluorescence and Whole Mounts analysis

Tissue biopsies were fixed in 4% neutral-buffered formalin (24h) and paraffin embedded. Paraffin-embedded tissues were sectioned and processed for immunohistochemistry (IHC) or hematoxylin and eosin (H/E) stainings. IHC was performed on 4μm sections using the following primary antibodies: anti-GFP (Thermofisher, A11122), anti-GP100 (Abcam, ab137078). Revelation was performed according to the manufacturer’s recommendations with secondary biotinylated anti-rabbit antibodies, streptavidin-peroxidase complex (ABC Vectastain kit, Vector Laboratories) and peroxidase substrate DAB (Vector Laboratories).

Immunofluorescence staining with anti-Sox10 (Abcam, ab155279) and anti-GFP (Abcam, ab13970) were performed on 4µm paraffin sections. Revelation was performed using Alexa594-conjugated anti-rabbit, Alexa594-conjugated anti-chicken secondary antibodies and Alexa488-coupled secondary antibodies respectively (Invitrogen).

For Whole Mounts, mice tail skins were peeled and cut into small pieces. Skin samples were incubated in 5 mM EDTA-PBS during 4 hours at 37ᵒC. Tail epidermis was separated from dermis and fixed in 4% Formaldehyde (Sigma) for 2 hours at room temperature. For immunostaining, samples were blocked and permeabilized in PB buffer containing 0,5% milk, 0,25% gelatin, 0,5% Triton X-100 in TBS (0,9% NaCI, 20 mM Hepes) for 30 minutes at room temperature, and then incubated with a goat anti-SOX10 (Santa Cruz, ref. sc17342) and a chicken anti-GFP antibodies (Invitrogen, ref. A10262) in PB buffer overnight at 4°C. Samples were washed with 0,2% Tween20 in PBS and incubated with a mix of Alexa 555 anti-goat and Alexa 488 anti-chicken (Thermofisher) and dapi in PB buffer overnight at 4°C and mounted on coverglass with Mowiol (Biovalley).

### In situ hybridization

4um thickness FFPE mouse skin sections were deparaffined in xylene solution followed by ethanol dehydratation. RNAscope was performed with probes and a kit following manufacturer’s instructions (Advanced Cell Diagnostics, Hayward, CA). This includes treatment with hydrogen peroxide (10 min at RT), Target Retrieval (15 min at 98-100°C) and Protease Plus treatment (30 min at RT), as provided with the kit. RNAscope Multiplex Fluorescent V2 assay was performed with probes against mouse *Mitf* (Mm-Mitf-C1, ref. 422501, ACD), mouse *E4f1* (Mm-E4f1-C3, ref. 825641-C3, ACD), mouse *Elp3* (Mm-Elp3-C3, ref. 400941-C3, ACD) and mouse *Eef1g* (Mm-Eef1g-O1-C1, ref. 1827341-C1, ACD). TSA Vivid dyes 650 (ref.323273, ACD) and 570 (ref. 323272, ACD) were respectively used at 1:1000 dilution to detect the probes. RNA quality was checked with a positive probe mix (ref.320881, ACD) against RNA polymerase II subunit A (PolR2A-C1) and Peptidylprolyl Isomerase B (PPIB-C2). Non-specific signal and background were detected with a negative control probe (ref.320871, ACD) that targets the bacterial gene dihydrodipicolinate B. subtilis reductase (DapB). Immediately after fluorescent RNA labeling, tissue sections were incubated 1h at room temperature with mouse FcR Blocking Reagent (ref.130-092-575, Miltenyi Biotec) diluted in SuperBlock solution (ref.37535, ThermoFisher) then incubated overnight at 4°C with a rabbit monoclonal anti-GFP antibody (CST, ref. 2956) diluted at 1:100 in 1% BSA. The day after, the tissue was gently washed in PBS/0,2% Triton X100 and then incubated with secondary antibody donkey anti-rabbit Alexa Fluor488 (ref. A21206, Invitrogen) diluted 1:500 and DAPI 1:1000 in 1% BSA for 30min. Finally, the slides were washed in PBS/0,2% Triton X100 and coverslipped with ProLong Glass mounting medium (P36930, Invitrogen). For microscopic imaging, tissue specimens were acquired using a Thunder epifluorescent microscope (Leica Microsystems CMS GmbH, Wetzlar (Germany)) with laser excitation wavelengths of 340-380nm, 450-490nm, 530-560nm and 590-650 nm, with the 40× objective. The number of *E4f1*, *Mitf, Elp3 and Eef1g* mRNA transcripts determined by RNAscope was based on the fluorescent spot count of around 60-100 GFP positive melanocytes in microscopic images.

### Cell Culture and treatment

Melan-a cells were provided by Pr Dorothy Bennett’s laboratory (SGUL) and cultured in RPMI medium complemented with 10% fetal bovine serum (FBS, Eurobio), 1% penicillin/streptomycin, 1% D-glutamine, 1µM tetradecanoyl phorbol acetate (TPA, Sigma) at 37°C in 5% CO2. For puromycin incorporation assays, cells were treated with puromycin (0,1ug/ml) for 30min before sample collection.

### Lentiviral Particle Production and Infections

Lentiviral particles were produced in 293T packaging cells by transient transfection, using Jet-PEI reagent (Ozyme), of pLKO vectors encoding either a control ShRNA (Sigma, SHC002), shRNAs targeting mouse *E4f1* (Sigma-Mission, TRCN0000321451 and TRCN0000321377), mouse *Elp3* (Sigma-Mission, TRCN0000039309, TRCN0000039312 and TRCN0000326972) or mouse sh*eEF1g* (Sigma-Mission, TRCN0000244499, TRCN0000245388). Seventy-two hours after transfection, viral supernatants were harvested, filtered on 0,4um and added on Melan-a cells for 8 hours in presence of polybrene (5 μg/mL, Sigma). Antibiotic selection of transduced Melan-a was performed 48h after transduction with puromycin (1 μg/mL; Invotrogen), neomycin (concentration).

### RNA extraction and quantitative PCR

Melan-a cells were lysed in TriReagent (Sigma) and total RNAs were isolated according to the manufacturer’s recommendations. cDNAs were synthesized from 500ng of total RNA using SuperScript™ III Reverse Transcriptase (Invitrogen). Quantitative real-time PCR (RT-qPCR) was performed on a LightCycler 480 apparatus (Roche). The relative mRNA copy number was calculated using Ct values and was normalized to *Rpl13a* and Tubulin b5 mRNA levels. Primers sequences used are listed in Supplemental Table S2.

### RNA sequencing and Transcriptome analyses

Libraries were sequenced using an Illumina NovaSeq 6000 at the iGenSeq facility (Institut du Cerveau, Paris), generating 766 million 101bp x 2 paired-end reads (42.6 million per sample in average). Raw sequencing reads were de-multiplexed with Illumina *CASAVA v.1.8.2*. After quality check using *FastQC v0.11.5*, reads were cleaned using *cutadapt v2.9* to remove Illumina TruSeq adapters, low quality bases and reads, and discard reads shorter than 25 bp after filtering. Fragments were then aligned to the mouse genome (mm10) using *STAR v2.7.3a* following ENCODE best practice.

The raw gene-centric count matrix was built using *Rsubread* v2.4.3 R package using default parameters and considering the inbuild NCBI mm10 annotation. The count matrix was further normalized to account for sequencing depth differences and to remove unwanted variation applying a nonlinear full-quantile normalization (*EDAseq 2.24.0*) and a RUVs debatching strategy (*RUVSeq v1.24.0*). Only genes with more than one count per million in at least one sample were kept. The obtained normalized count matrix was used as an input for the differential expression analyses performed with *edgeR v3.32.1*, considering the treatment (control or shRNA against E4f1) and the surrogates in the design matrix and applying a Benjamini-Hocheberg correction for multiple-testing. Statistical significance was reached when the corrected *p*-value was smaller than 0.01, and the absolute fold-change greater than log_2_(1.5). The list of differentially expressed genes was used to perform a gene set over-representation analysis considering the Molecular Signature Database (*MSigDB v7.2*) Reactome gene sets using a right-tail modified Fisher’s exact test and the hypergeometric distribution to provide *p*-value. Gene Set Enrichment Analyses were conducted using the R package *fgsea v1.16.0* with 1000 permutations to obtain *p*-values.

### Protein extraction and western blotting

For total protein extracts, Melan-a cells were lysed in Laemmli buffer (80mM Tris pH=6,8, 2% SDS, 12% sucrose, 2% β-mercaptoethanol, bromophenol blue) supplemented with phosphatase (Phosphatase Inhibitor Cocktail 2, Sigma) and protease (cOmplete™ Protease Inhibitor Cocktail, Roche) inhibitors according to the manufacturer’s recommendations. Immunoblotting was performed using the following antibodies: anti-E4F1 (Santa Cruz Biotechnology, sc514718), anti-DLAT (Santa Cruz Biotechnology, sc-271534), anti-MITF (Sigma, HPA003259), anti-DCT (Abcam, ab74073), anti-ELP1 (AnaSpec, AS-54494), anti-ELP3 (Abcam, ab190907) and ACTIN (Sigma, A3854), eEF1g (Santa Cruz, sc-393378), eIF4b (Santa Cruz, sc-376062).

### Polysome profiling

Melan-a cells were lysed in polysome lysis buffer (7% sucrose, 50 mM Tris-HCl pH 7.5, 5 mM MgCl2, 25 mM KCl, 1% NP40, 10 U/mL RNAseOUT (Invitrogen, 10777019) supplemented with protease inhibitors and 20 μg/mL Emetin (Sigma, E2375)) by hard shaking with 1.4 mm ceramic spheres (Lysing matrix D MPBio, 1169130-CF) in a FastPrep instrument (MPBio). 30 μg of total RNA were loaded on 15–50% sucrose gradient and ultracentrifuged at 2,10,000×g for 2.5h at 4 °C in a SW41 rotor (Beckman Coulter). Approximately 10% of the cytosolic RNAs were kept as an input. The different fractions were separated through a live optical density (OD) 254 nm UV spectrometer and collected with an ISCO (Lincoln, NE) density gradient fractionation system. The absorbance at 254 nm was measured continuously as a function of gradient depth. Total cytosolic mRNAs, as well as mRNAs associated with 1 to 3 ribosomes (Light polysomal fraction) and mRNAs associated with more than 3 ribosomes (Heavy polysomal fraction) were extracted using TRIzol LS (Invitrogen, 10296028) according to the manufacturer instructions. For sequencing, mRNA libraries were generated following the manufacturer recommendations (mRNA hyper prep from ROCHE). Bar-coded libraries were pooled and sequenced on a Novaseq6000 200 cycles Illumina sequencer, with a sequencing depth corresponding to a minimum of 80 millions of 2 × 100 bp paired-end reads per sample after demultiplexing. Polysome profiling datasets were pre-processed using the nf-core RNA-seq pipeline v3.11.044. Initial quality control of raw reads was conducted with FastQC v0.11.9. Trim Galore v0.6.7 was used for reads trimming, with parameters set to a minimum read length of 25 bp, a quality threshold of Phred score 30, and a stringency of 10. Additionally, 1 bp was clipped from both 5’ ends of R1 and R2 reads. Reads were aligned to the mouse reference genome (GRCm38) using STAR v2.7.9a. Transcript-level quantification was performed using Salmon v1.10.1, and gene-level summarization was carried out using tximport v1.32.0. Post-alignment quality was assessed with RSeQC v3.0.1 to ensure data integrity and alignment accuracy. For statistical analysis, datasets from polysome fractions (light, heavy, and total) were analyzed independently using R v4.4.0. Differential expression (DE) analysis for each fraction was conducted with edgeR v4.2.1, comparing gene expression between *E4f1^(Tyr)KO^* and CTL animals. Normalized count data were fitted to a generalized linear model (GLM) according to E4f1 status as a cofactor (KO or WT). After estimating the signal dispersion, the abundance of transcripts in a given fraction was considered significantly different in likelihood ratio tests when exhibiting an absolute fold-change ≥ 1.2 and an FDR-adjusted p-value < 0.2. To integrate results from the different fractions, the corresponding genes were categorized as controlled at the posttranscriptional level (differential abundance of their transcripts in either the heavy and/or the light polysomal fractions, but not in the total fraction), at the transcriptional level (differential abundance of their transcripts in the total fraction but not in the heavy nor in the light polysomal fractions), and at both the transcriptional and posttrancriptional (Mix) levels (differential abundance of their transcripts in the total fraction and in the heavy and/or in the light polysomal fractions). Codon usage bias was assessed by comparing codon frequencies in transcripts controlled at the post-trancriptional level in E4F1-deficient cells to those in transcripts from the entire GRCm38 transcriptome using χ2 tests. In the corresponding proteins, enrichment of the specific hydrophilic motif linked to U34-dependent translational defects was conducted using the AME (Analysis of Motif Enrichment)45 tool from the MEME suite v1.12.0, using the occurrence of this motif in the full GRCm38 proteome as a reference.

### Statistical analysis

Statistical analyses were performed using GraphPad Prism (vs 11.1.0) and R. Fold-change data were log-transformed and analyzed using two-tailed one-sample t-tests against 0. Wilcoxon signed-rank tests were used where appropriate. Results are presented as geometric means with 95% confidence intervals. P values were adjusted for multiple comparisons using the Holm method. Adjusted P values < 0.05 were considered statistically significant.

## Acknowledgements

This work was supported by grants from the Agence Nationale pour la Recherche (ANR, METABOSKIN, JCJC ANR-17-CE13-0017), the Institut National du Cancer (INCa, PyrE4F1), the ARC foundation, the Ligue contre le cancer (Labellisation, Ligue régionale), GEFLUC (Occitanie) and with the institutional support of the Institut National de la santé et de la Recherche Médicale (INSERM). We are grateful for the technical support of the animal facilities of Montpellier Biocampus. We acknowledge the imaging facility MRI, member of the national infrastructure France-BioImaging (https://ror.org/01y7vt929) supported by the French National Research Agency (ANR-24-INBS-0005 FBI BIOGEN)”. The RHEM (Réseau d’Histologie Expérimentale de Montpellier) histology facility was supported by SIRIC Montpellier Cancer (Grant INCa_Inserm_DGOS_12553, the European Regional Development Foundation, and the Occitanie region (FEDER-FSE 2014-2020 Languedoc Roussillon). We thank members of the LLC team for critical reading of the manuscript.

## Author Contributions

RM, HM, HL, SB, PL, CC, LCL and LM designed the experiments and analyzed, discussed, and interpreted the data. RM, HM, HL, SB, EB, DP, FC, RL and TA performed the in vivo experiments. RM, HM, HL, SB, PL, GP, SA and PL performed the in vitro experiments. RM, HM, DMM, CA, RPF and DA performed and analyzed the polysome profiling. FPC, EB, DP, FC, PN, BF and LL performed the histological analyses. RM, HM, LCL and LM wrote the paper. All authors gave final approval of the submitted manuscript.

---

**Figure S1:**
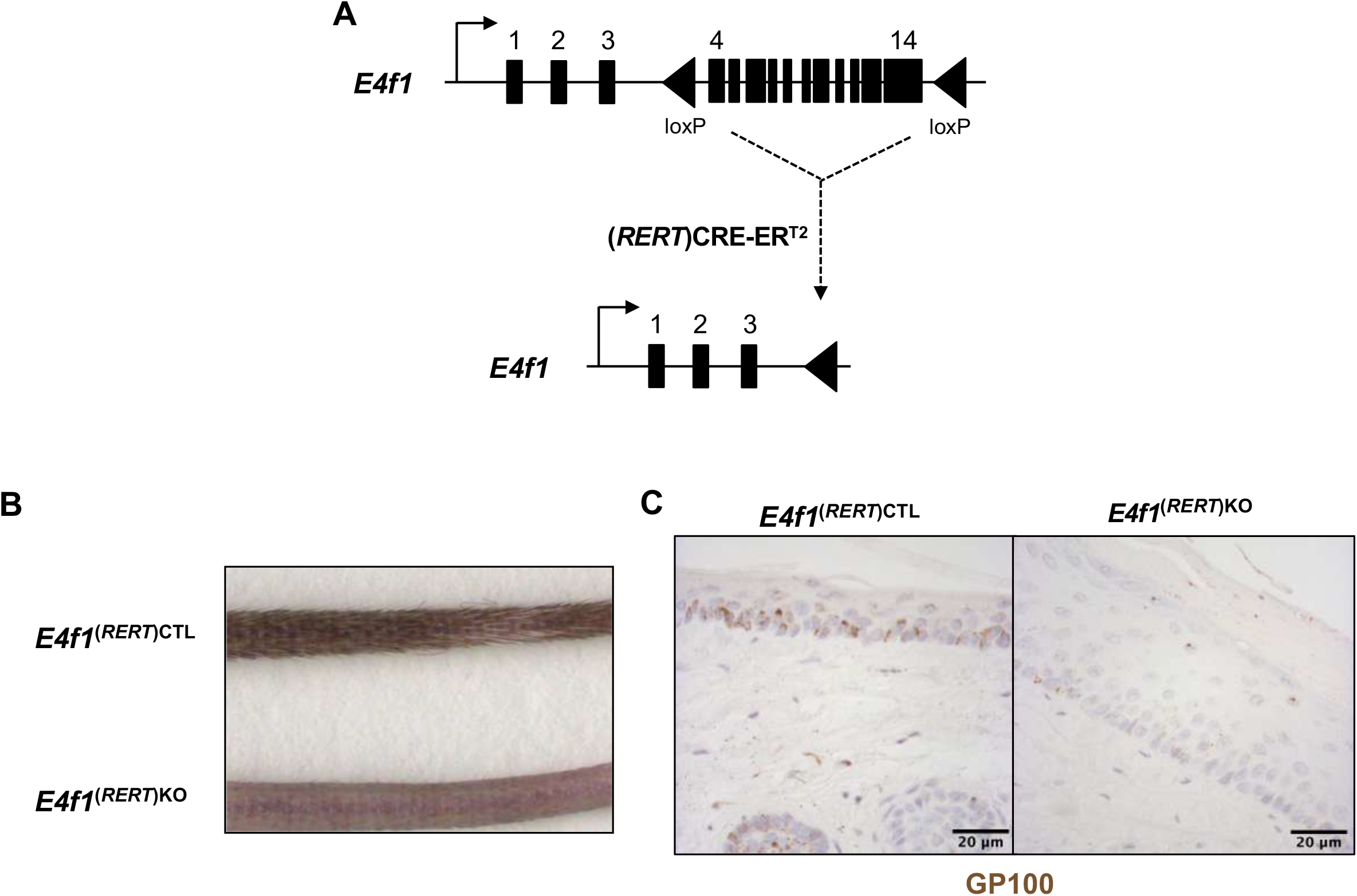
E4f1 whole-body conditional knock-out induces skin pigmentation defects. **(A)** Schematic representation of the *E4f1*^flox^ allele before/after (RERT)CreER^T2^ recombination. **(B)** Macroscopic observation of the tail of *E4f1*^(*RERT*)CTL^ and *E4f1*^(*RERT*)KO^ mice. *E4f1*^(*RERT*)KO^ animals display skin pigmentation defects. **(C)** Immunohistochemistry analysis of the GP100 melanosome marker on tail skin sections prepared from *E4f1*^(*RERT*)CTL^ and *E4f1*^(*RERT*)KO^ mice. Scale bar, 20 µm.

**Figure S2:**
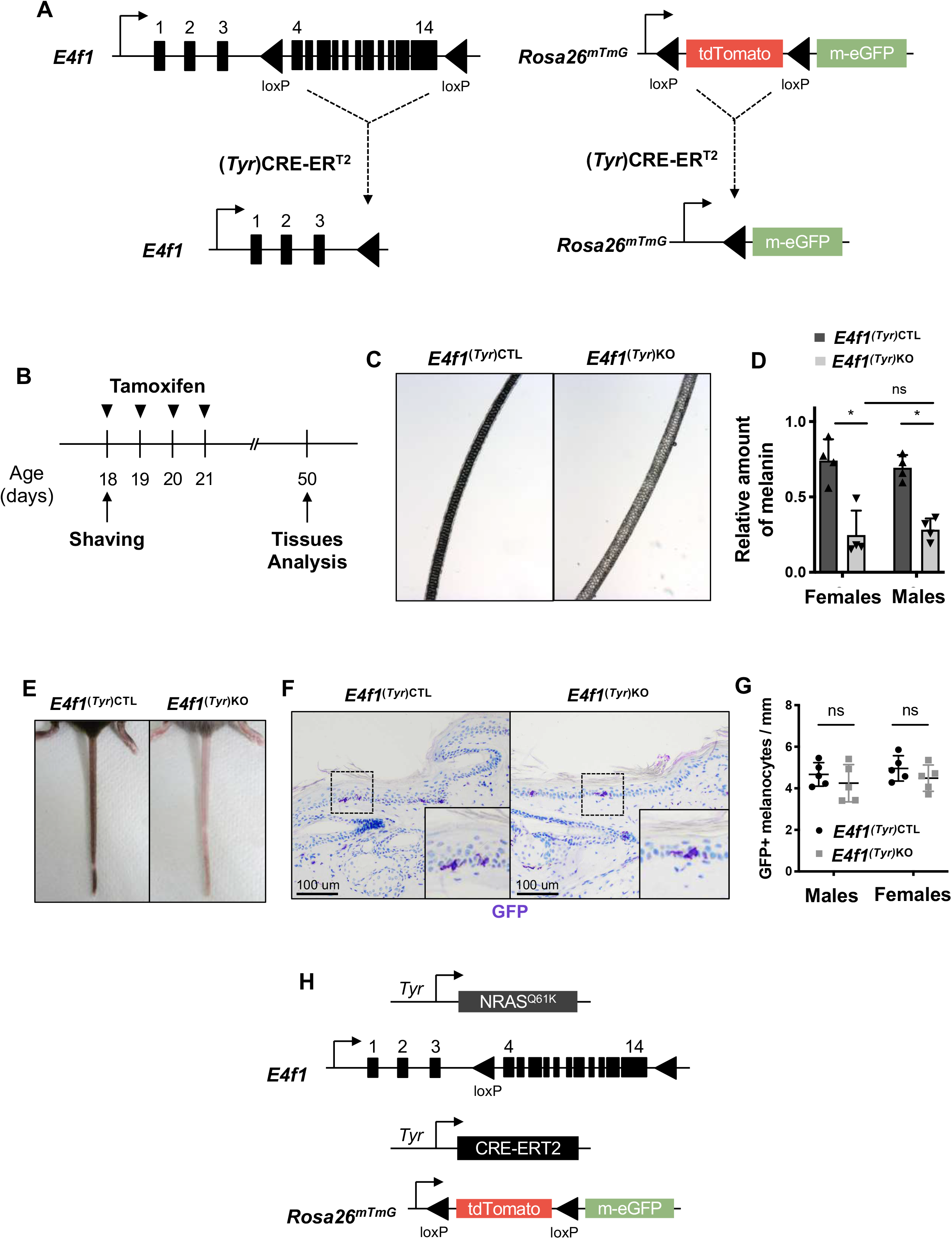
Conditional *E4f1* inactivation in melanocytes. **(A)** Schematic representation of the *E4f1*^flox^ and mTmG alleles before/after (Tyr)CreER^T2^ recombination. **(B)** Experimental procedure of tamoxifen administration to *E4f1*^(*Tyr*)CTL^ and *E4f1*^(*Tyr*)KO^ mice. 18 days-old mice are shaved and treated with topical application of 2 mg of tamoxifen on back and tail skin (1 application per day for 4 days). Tissues are analyzed one month after treatment (day 50). (**C**) Microscopic observation of hairs from *E4f1^(Tyr)CTL^*et *E4f1*^(*Tyr*)KO^ animals. (**D**) Melanin quantification in hairs from *E4f1*^(*Tyr*)CTL^ et *E4f1*^(*Tyr*)KO^ females and males (mean ± SEM, n=4). (**E**) Macroscopic observation of *E4f1^(Tyr)CTL^* et *E4f1^(Tyr^*^)*KO*^ tail. *E4f1^(Tyr)KO^* animals exhibits skin pigmentation defects compared to control littermates. **(F)** Immunohistochemistry analysis of the GFP reporter on tail skin sections prepared from *E4f1^(Tyr^*^)CTL^ et *E4f1*^(*Tyr*)KO^ mice and showing inter follicular melanocytes in the epidermis (arrow). Insets show images at higher magnification. Scale bar, 100 µm. **(G)** Quantification of GFP^+^ melanocytes per millimeter of tail skin from males and females of *E4f1*^(*Tyr*)CTL^ et *E4f1*^(*Tyr*)KO^ mice (mean ± SEM, n=5 with 100 cells analyzed in each condition). (**H**) Schematic representation of genetic constructs allowing *Nras^Q61K^* constitutive expression in melanocytes, *E4f1* inactivation in melanocytes upon tamoxifen-induced TyrCreER^T2^ recombination, and mTmG reporter expression.

**Figure S3:**
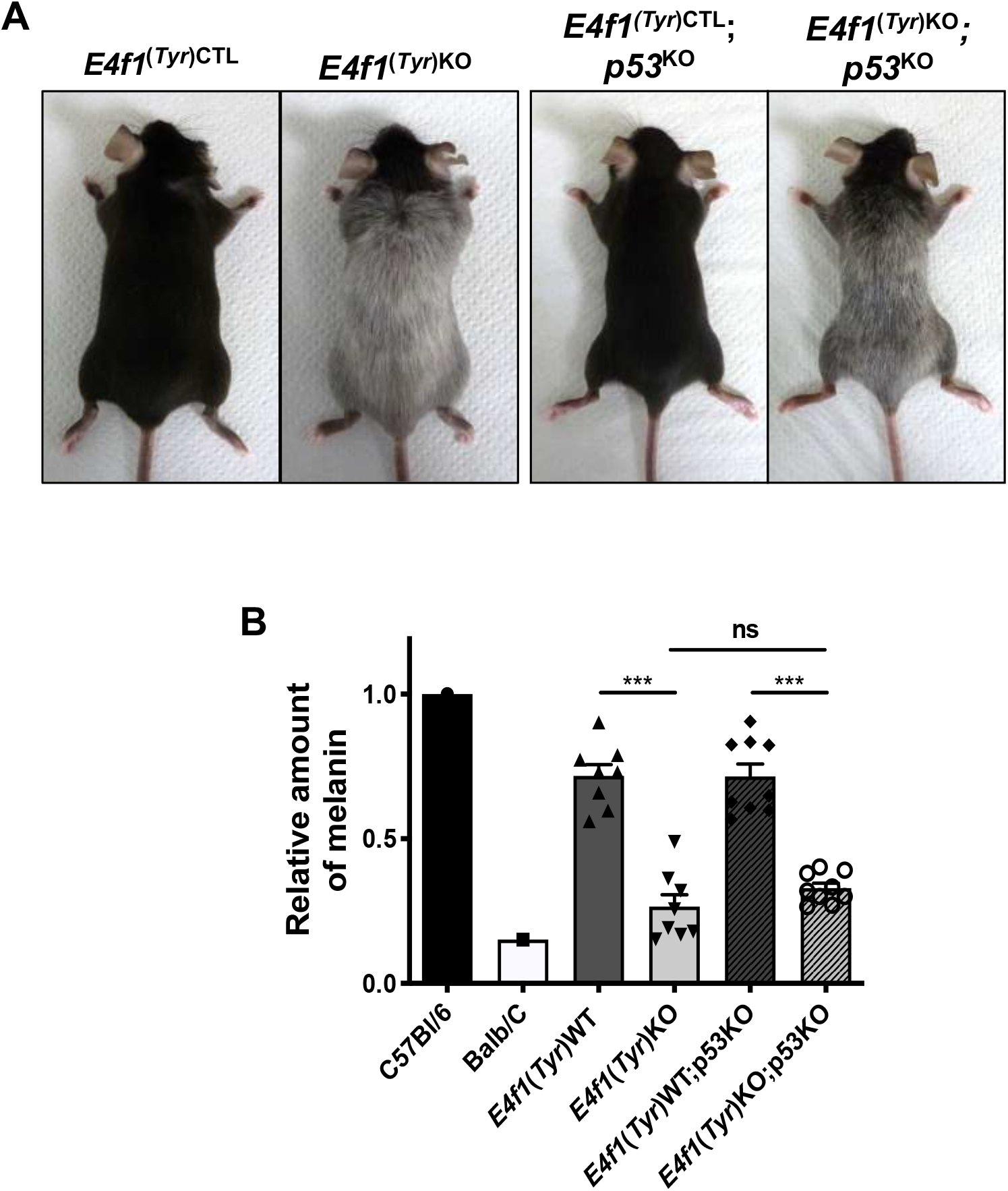
*E4f1*-mediated pigmentation defects are not p53-dependent. **(A)** Macroscopic observation of *E4f1*^(*Tyr*)CTL^; p53^KO^ and *E4f1*^(*Tyr*)KO^; p53^KO^ mice, compared to their *E4f1*^(*Tyr*)CTL^ and *E4f1*^(*Tyr*)KO^ littermates **(B)** Melanin quantification in hairs from *E4f1*^(*Tyr*)CTL^; p53^KO^ and *E4f1*^(*Tyr*)KO^; p53^KO^ mice, compared to their *E4f1*^(*Tyr*)CTL^ and *E4f1*^(*Tyr*)KO^ littermates (mean ± SEM, n=8). Hairs from C57Bl/6 and Balb/C mice were used as positive and negative control, respectively.

**Figure S4:**
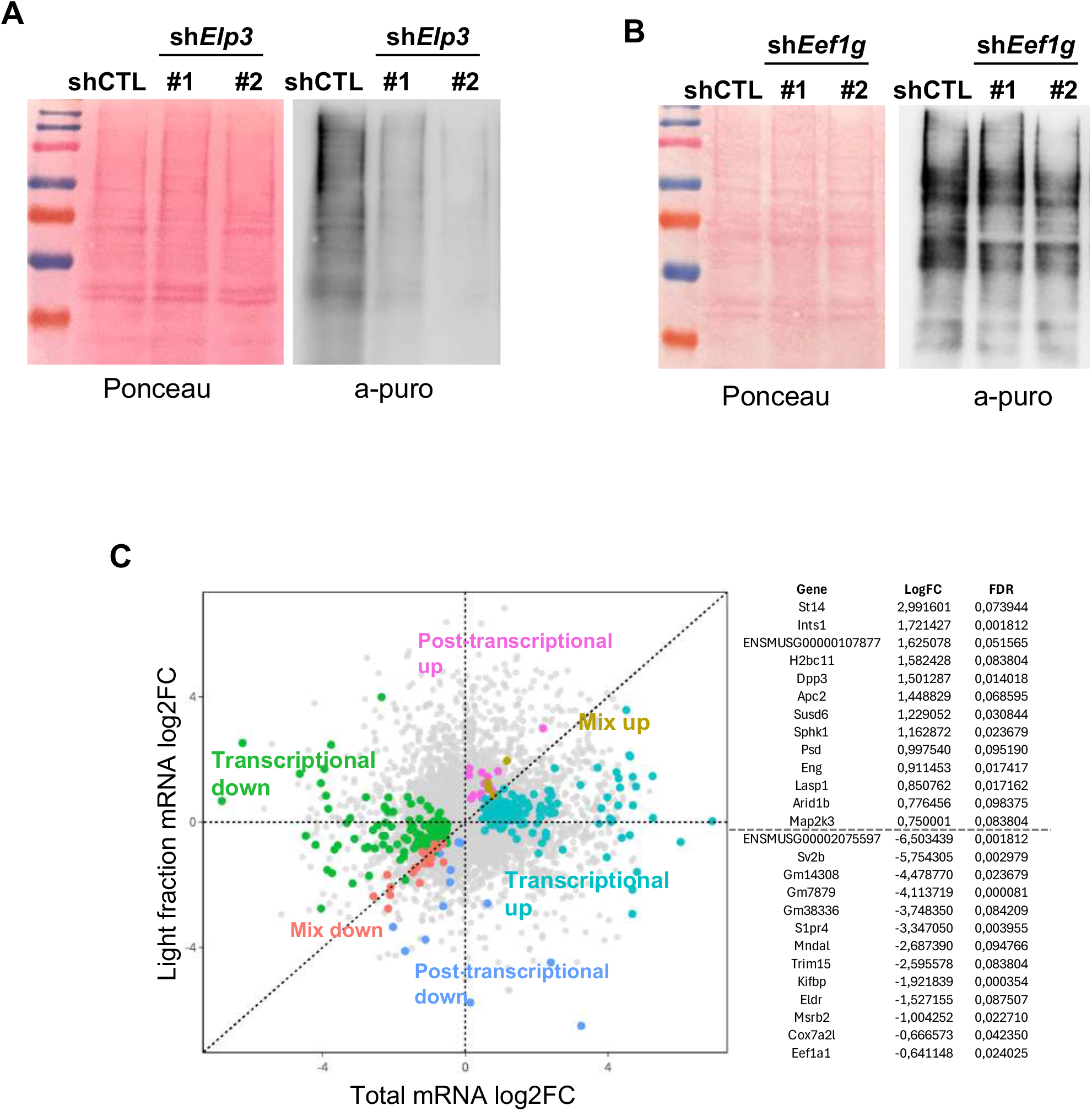
(**A**) Immunoblot analysis of puromycin in Melan-a cells, 10 days after transduction with lentiviral particles encoding an irrelevant shRNA (shCTL) or shRNA targeting *Elp3* (sh*Elp3*). (**B**) Immunoblot analysis of puromycin in Melan-a cells, 10 days after transduction with lentiviral particles encoding an irrelevant shRNA (shCTL) or shRNA targeting *Eef1g* (sh*Eef1g*). (**C**) Polysome profiling analysis of brains of shCTL and sh*E4f1* Melan-a cells (n=5). Dot plots represent transcript levels (Log_2_ fold change) between sh*E4f1* and shCTL Melan-a cells in the pooled light (less than 3 ribosomes) polysomal fractions, according to their abundance in the total cytosolic fraction. The lists indicate the 16 genes which transcript abundance is the most deregulated (log_2_FC) in the heavy polysomal fractions prepared from sh*E4f1* Melan-a cells but showing similar total transcript levels.

**Figure S5:**
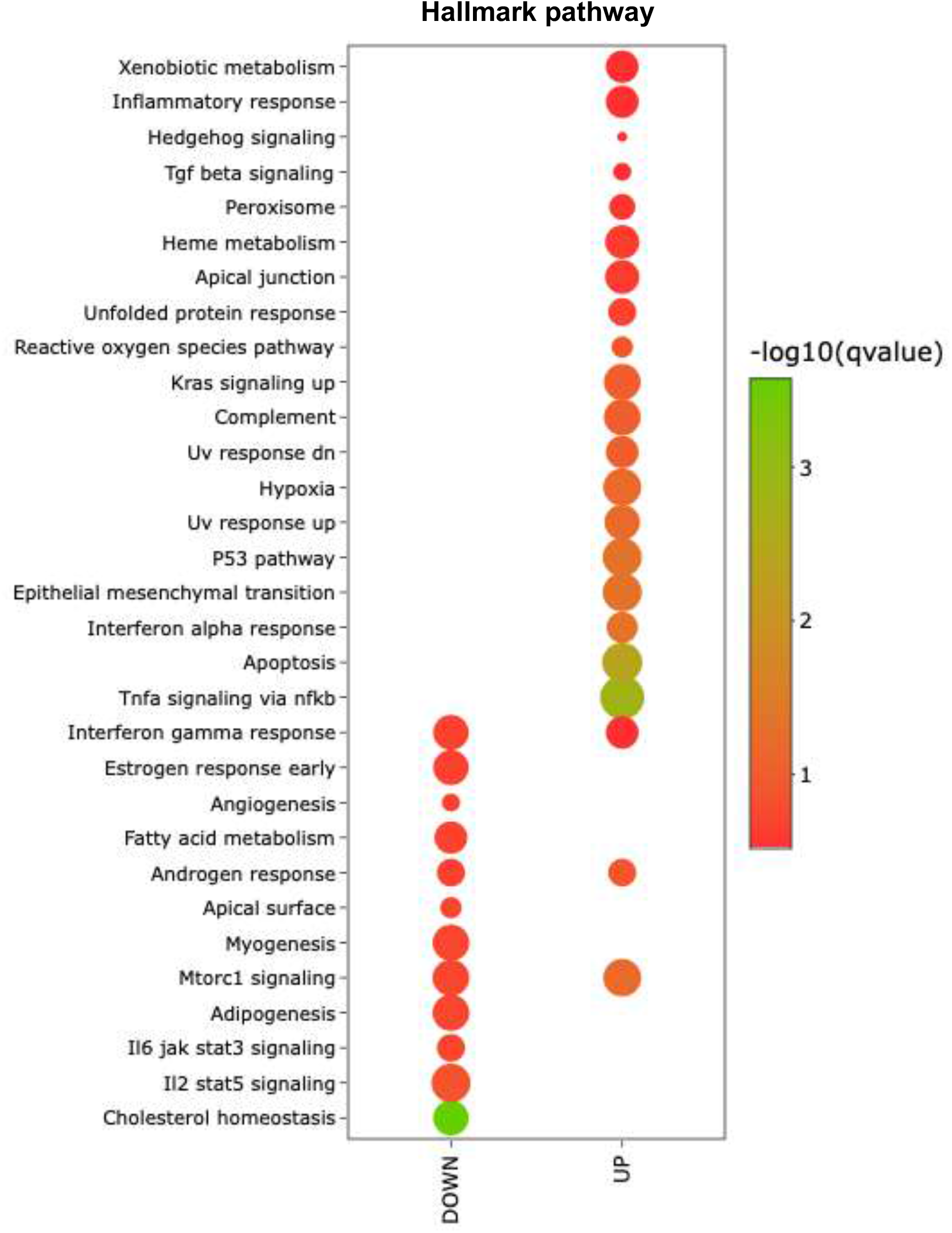
Hallmark pathways up and down regulated in Sh*E4f1* Melan-a cells.

**Supplemental Table S1:**
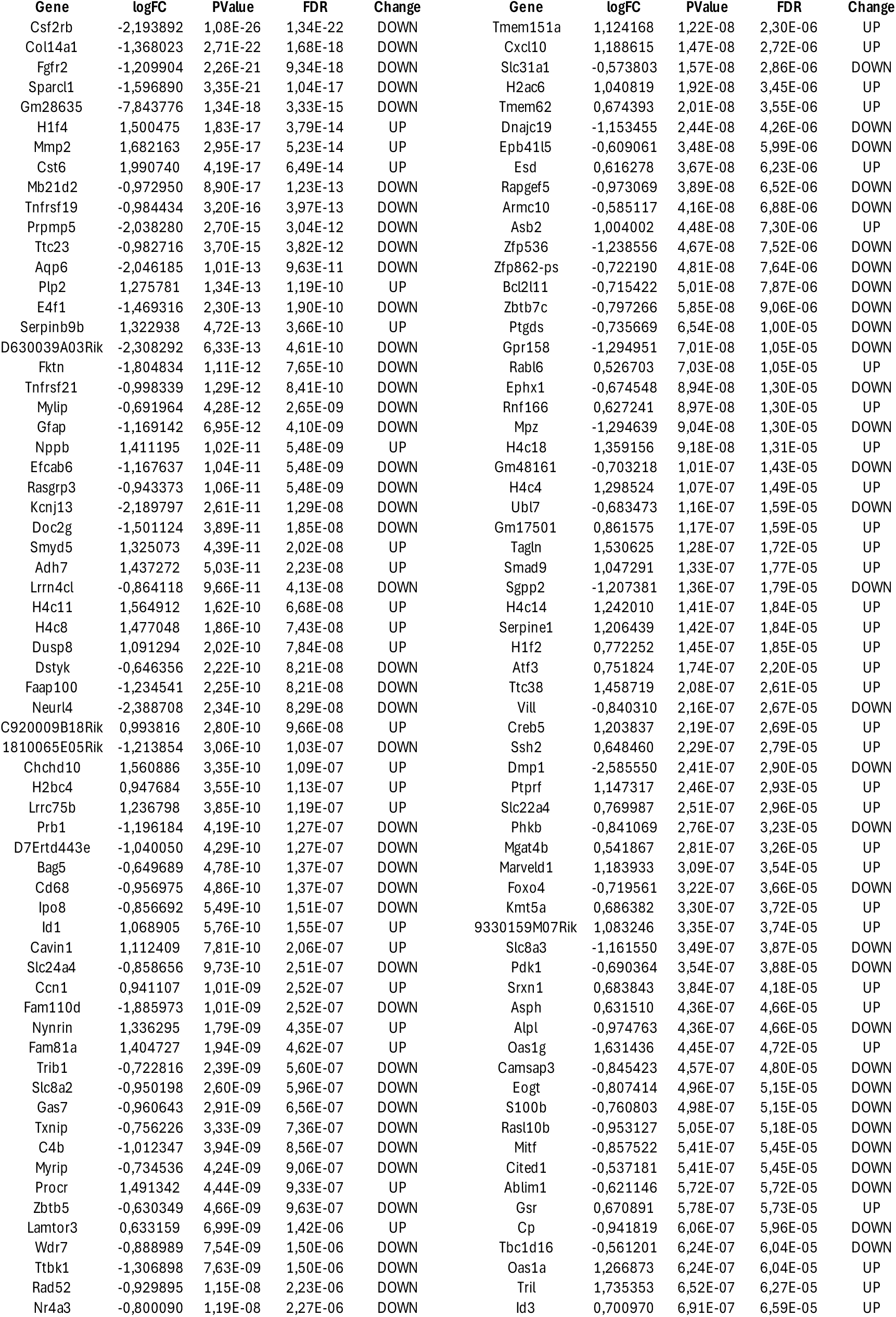
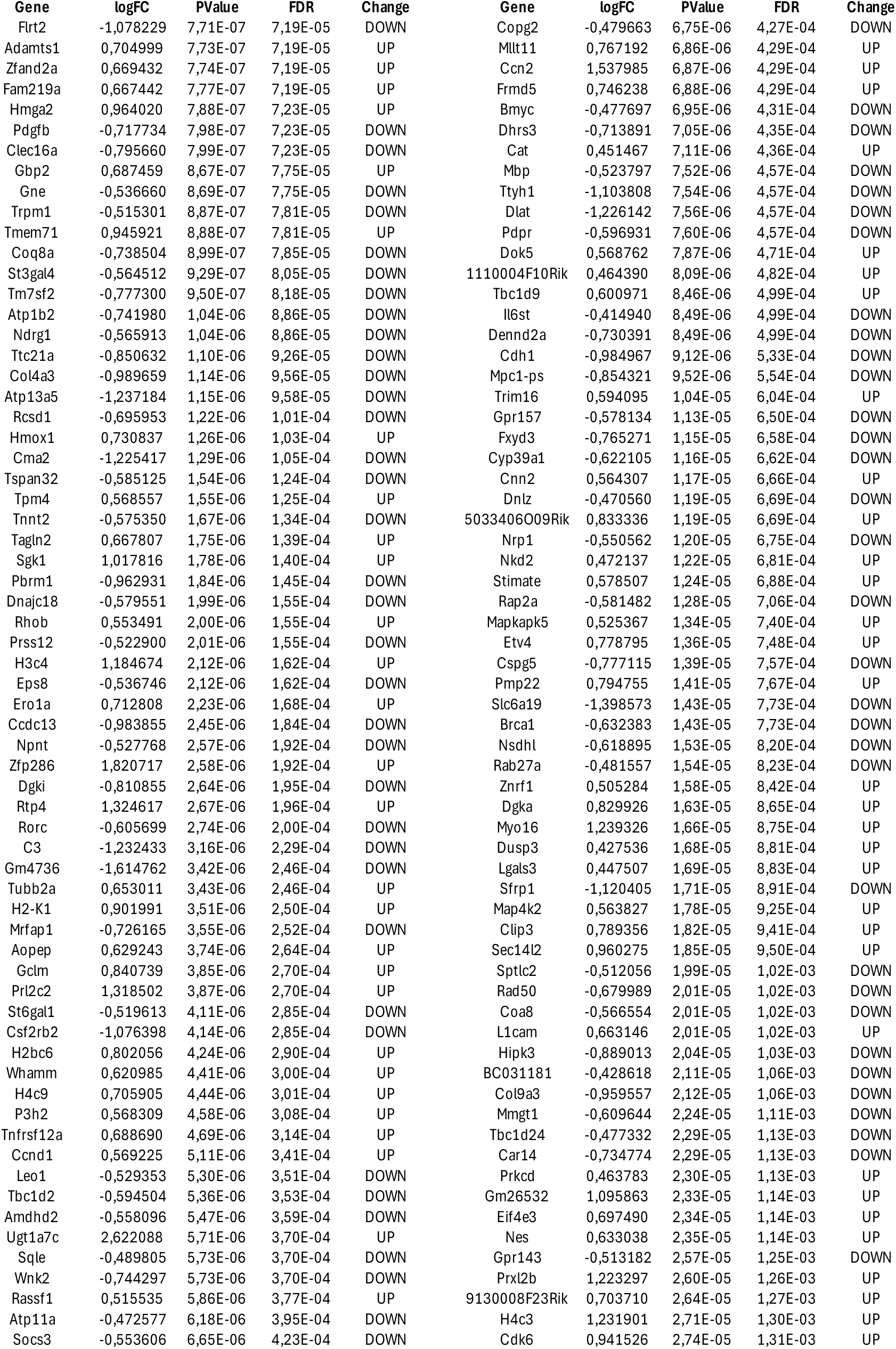
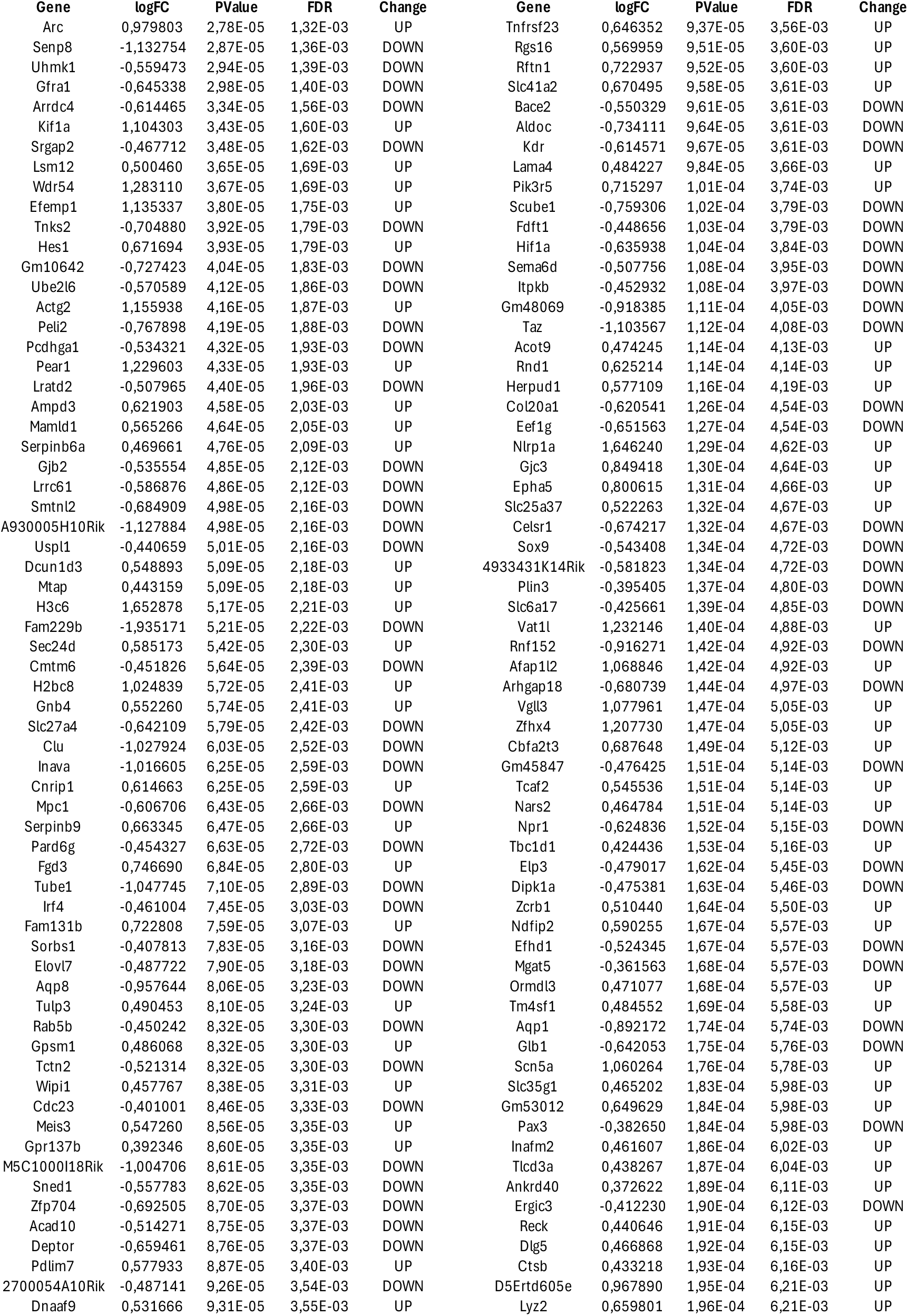
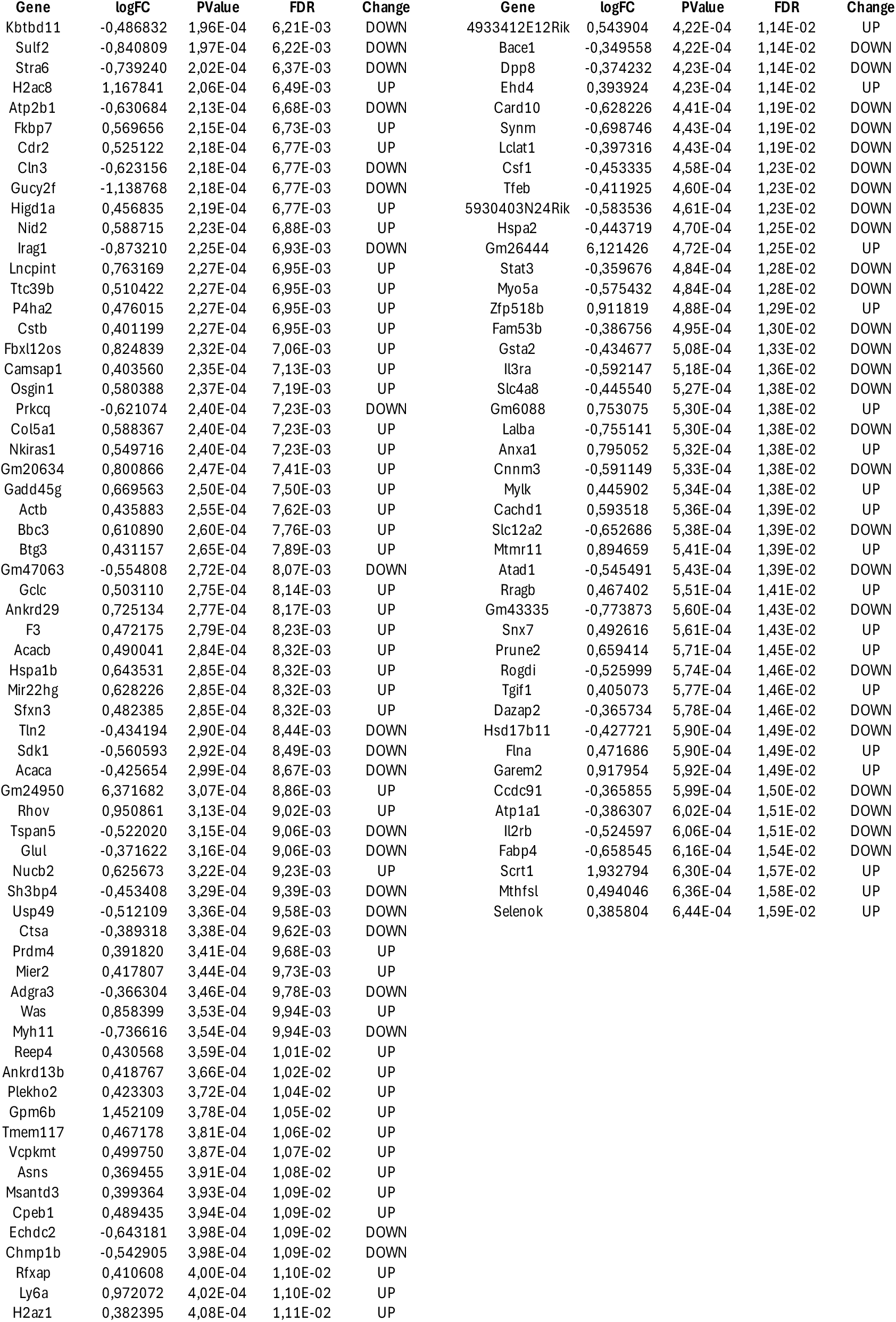
Top 500 more significant deregulated genes upon E4f1 inactivation in Melan-a cells.

**Supplemental Table S2:**
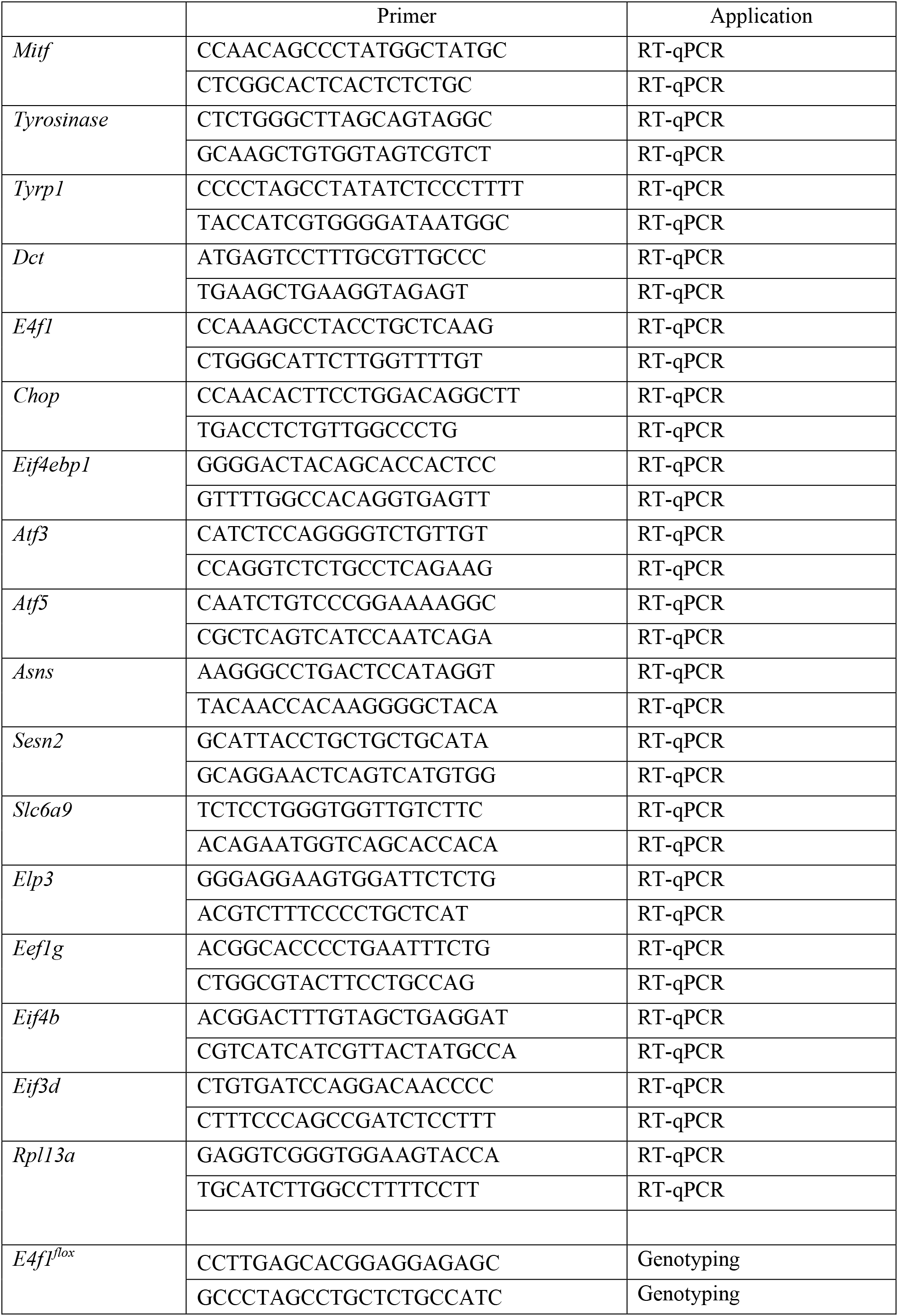
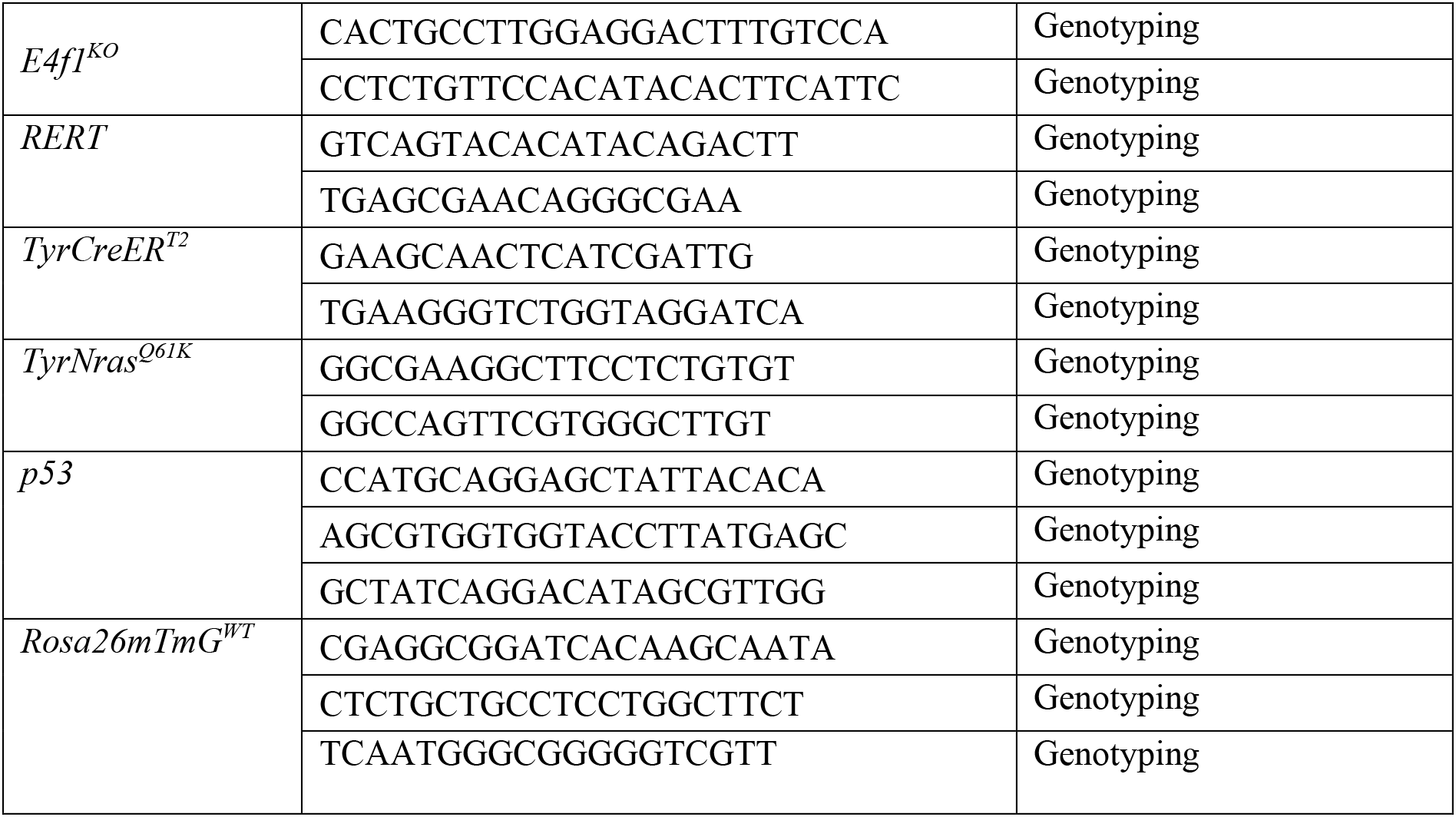
Primers used for RT-qPCR and mouse genotyping.

## REFERENCES

1. Kobayashi T, Imokawa G, Bennett DC, Hearing VJ. Tyrosinase stabilization by Tyrp1 (the brown locus protein). J Biol Chem. 1998;273: 31801–31805. doi:10.1074/jbc.273.48.31801

2. Meredith P, Sarna T. The physical and chemical properties of eumelanin. Pigment Cell Res. 2006;19: 572–594. doi:10.1111/j.1600-0749.2006.00345.x

3. Ando H, Niki Y, Ito M, Akiyama K, Matsui MS, Yarosh DB, et al. Melanosomes are transferred from melanocytes to keratinocytes through the processes of packaging, release, uptake, and dispersion. J Invest Dermatol. 2012;132: 1222–1229. doi:10.1038/jid.2011.413

4. Yamaguchi Y, Brenner M, Hearing VJ. The Regulation of Skin Pigmentation *. J Biol Chem. 2007;282: 27557–27561. doi:10.1074/jbc.R700026200

5. Kondo T, Hearing VJ. Update on the regulation of mammalian melanocyte function and skin pigmentation. Expert Rev Dermatol. 2011;6: 97–108. doi:10.1586/edm.10.70

6. del Marmol V, Beermann F. Tyrosinase and related proteins in mammalian pigmentation. FEBS Lett. 1996;381: 165–168. doi:10.1016/0014-5793(96)00109-3

7. Bentley NJ, Eisen T, Goding CR. Melanocyte-specific expression of the human tyrosinase promoter: activation by the microphthalmia gene product and role of the initiator. Mol Cell Biol. 1994;14: 7996–8006. doi:10.1128/mcb.14.12.7996-8006.1994

8. Yasumoto K, Yokoyama K, Shibata K, Tomita Y, Shibahara S. Microphthalmia-associated transcription factor as a regulator for melanocyte-specific transcription of the human tyrosinase gene. Mol Cell Biol. 1994;14: 8058–8070. doi:10.1128/mcb.14.12.8058-8070.1994

9. Yasumoto K, Yokoyama K, Takahashi K, Tomita Y, Shibahara S. Functional analysis of microphthalmia-associated transcription factor in pigment cell-specific transcription of the human tyrosinase family genes. J Biol Chem. 1997;272: 503–509. doi:10.1074/jbc.272.1.503

10. Goding CR, Arnheiter H. MITF—the first 25 years. Genes Dev. 2019;33: 983–1007. doi:10.1101/gad.324657.119

11. Hertwig P. Neue Mutationen und Koppelungsgruppen bei der Hausmaus. Z Für Indukt Abstamm-Vererbungslehre. 1942;80: 220–246. doi:10.1007/BF01741984

12. Steingrímsson E, Copeland NG, Jenkins NA. Melanocytes and the microphthalmia transcription factor network. Annu Rev Genet. 2004;38: 365–411. doi:10.1146/annurev.genet.38.072902.092717

13. Léger S, Balguerie X, Goldenberg A, Drouin-Garraud V, Cabot A, Amstutz-Montadert I, et al. Novel and recurrent non-truncating mutations of the MITF basic domain: genotypic and phenotypic variations in Waardenburg and Tietz syndromes. Eur J Hum Genet EJHG. 2012;20: 584–587. doi:10.1038/ejhg.2011.234

14. Pingault V, Ente D, Dastot-Le Moal F, Goossens M, Marlin S, Bondurand N. Review and update of mutations causing Waardenburg syndrome. Hum Mutat. 2010;31: 391–406. doi:10.1002/humu.21211

15. Amiel J, Watkin PM, Tassabehji M, Read AP, Winter RM. Mutation of the MITF gene in albinism-deafness syndrome (Tietz syndrome). Clin Dysmorphol. 1998;7: 17–20.

16. Smith SD, Kelley PM, Kenyon JB, Hoover D. Tietz syndrome (hypopigmentation/deafness) caused by mutation of MITF. J Med Genet. 2000;37: 446–448. doi:10.1136/jmg.37.6.446

17. Seberg HE, Van Otterloo E, Cornell RA. Beyond MITF: Multiple transcription factors directly regulate the cellular phenotype in melanocytes and melanoma. Pigment Cell Melanoma Res. 2017;30: 454–466. doi:10.1111/pcmr.12611

18. Hemesath TJ, Price ER, Takemoto C, Badalian T, Fisher DE. MAP kinase links the transcription factor Microphthalmia to c-Kit signalling in melanocytes. Nature. 1998;391: 298–301. doi:10.1038/34681

19. Wang C, Zhao L, Su Q, Fan X, Wang Y, Gao S, et al. Phosphorylation of MITF by AKT affects its downstream targets and causes TP53-dependent cell senescence. Int J Biochem Cell Biol. 2016;80: 132–142. doi:10.1016/j.biocel.2016.09.029

20. Takeda K, Takemoto C, Kobayashi I, Watanabe A, Nobukuni Y, Fisher DE, et al. Ser298 of MITF, a mutation site in Waardenburg syndrome type 2, is a phosphorylation site with functional significance. Hum Mol Genet. 2000;9: 125–132. doi:10.1093/hmg/9.1.125

21. Xu W, Gong L, Haddad MM, Bischof O, Campisi J, Yeh ETH, et al. Regulation of Microphthalmia-Associated Transcription Factor MITF Protein Levels by Association with the Ubiquitin-Conjugating Enzyme hUBC9. Exp Cell Res. 2000;255: 135–143. doi:10.1006/excr.2000.4803

22. Murakami H, Arnheiter H. Sumoylation modulates transcriptional activity of MITF in a promoter-specific manner. Pigment Cell Res. 2005;18: 265–277. doi:10.1111/j.1600-0749.2005.00234.x

23. Falletta P, Sanchez-del-Campo L, Chauhan J, Effern M, Kenyon A, Kershaw CJ, et al. Translation reprogramming is an evolutionarily conserved driver of phenotypic plasticity and therapeutic resistance in melanoma. Genes Dev. 2017;31: 18–33. doi:10.1101/gad.290940.116

24. Raychaudhuri P, Rooney R, Nevins JR. Identification of an E1A-inducible cellular factor that interacts with regulatory sequences within the adenovirus E4 promoter. EMBO J. 1987;6: 4073–4081.

25. Lee KA, Green MR. A cellular transcription factor E4F1 interacts with an E1a-inducible enhancer and mediates constitutive enhancer function in vitro. EMBO J. 1987;6: 1345–1353.

26. Le Cam L, Linares LK, Paul C, Julien E, Lacroix M, Hatchi E, et al. E4F1 Is an Atypical Ubiquitin Ligase that Modulates p53 Effector Functions Independently of Degradation. Cell. 2006;127: 775–788. doi:10.1016/j.cell.2006.09.031

27. Rizos H, Diefenbach E, Badhwar P, Woodruff S, Becker TM, Rooney RJ, et al. Association of p14ARF with the p120E4F transcriptional repressor enhances cell cycle inhibition. J Biol Chem. 2003;278: 4981–4989. doi:10.1074/jbc.M210978200

28. Chagraoui J, Niessen SL, Lessard J, Girard S, Coulombe P, Sauvageau M, et al. E4F1: a novel candidate factor for mediating BMI1 function in primitive hematopoietic cells. Genes Dev. 2006;20: 2110–2120. doi:10.1101/gad.1453406

29. Rodier G, Kirsh O, Baraibar M, Houlès T, Lacroix M, Delpech H, et al. The Transcription Factor E4F1 Coordinates CHK1-Dependent Checkpoint and Mitochondrial Functions. Cell Rep. 2015;11: 220–233. doi:10.1016/j.celrep.2015.03.024

30. Lacroix M, Rodier G, Kirsh O, Houles T, Delpech H, Seyran B, et al. E4F1 controls a transcriptional program essential for pyruvate dehydrogenase activity. Proc Natl Acad Sci U S A. 2016;113: 10998–11003. doi:10.1073/pnas.1602754113

31. Lacroix M, Linares LK, Rueda-Rincon N, Bloch K, Di Michele M, De Blasio C, et al. The multifunctional protein E4F1 links p53 to lipid metabolism in adipocytes. Press.

32. Di Michele M, Laguesse S, Attina A, Mathieu H, Rousseau M, Jahanault-Tagliani C, et al. E4f1 coordinates pyruvate metabolism and the activity of the Elongator complex to ensure protein translation fidelity during neuronal development. Prep.

33. Le Cam L, Lacroix M, Ciemerych MA, Sardet C, Sicinski P. The E4F Protein Is Required for Mitotic Progression during Embryonic Cell Cycles. Mol Cell Biol. 2004;24: 6467–6475. doi:10.1128/MCB.24.14.6467-6475.2004

34. Lacroix M, Caramel J, Goguet-Rubio P, Linares LK, Estrach S, Hatchi E, et al. Transcription factor E4F1 is essential for epidermal stem cell maintenance and skin homeostasis. Proc Natl Acad Sci U S A. 2010;107: 21076–21081. doi:10.1073/pnas.1010167107

35. Goguet-Rubio P, Seyran B, Gayte L, Bernex F, Sutter A, Delpech H, et al. E4F1-mediated control of pyruvate dehydrogenase activity is essential for skin homeostasis. Proc Natl Acad Sci U S A. 2016;113: 11004–11009. doi:10.1073/pnas.1602751113

36. Caramel J, Lacroix M. E4F1 connects the Bmi1-ARF-p53 pathway to epidermal stem cell-dependent skin homeostasis. Cell Cycle.: 3.

37. Cui R, Widlund HR, Feige E, Lin JY, Wilensky DL, Igras VE, et al. Central Role of p53 in the Suntan Response and Pathologic Hyperpigmentation. Cell. 2007;128: 853–864. doi:10.1016/j.cell.2006.12.045

38. Vivas-García Y, Falletta P, Liebing J, Louphrasitthiphol P, Feng Y, Chauhan J, et al. Lineage-Restricted Regulation of SCD and Fatty Acid Saturation by MITF Controls Melanoma Phenotypic Plasticity. Mol Cell. 2020;77: 120–137.e9. doi:10.1016/j.molcel.2019.10.014

39. Hatchi E, Rodier G, Lacroix M, Caramel J, Kirsh O, Jacquet C, et al. E4F1 deficiency results in oxidative stress–mediated cell death of leukemic cells. J Exp Med. 2011;208: 1403–1417. doi:10.1084/jem.20101995

40. Hatchi E, Rodier G, Sardet C, Le Cam L. E4F1 dysfunction results in autophagic cell death in myeloid leukemic cells. Autophagy. 2011;7: 1566–1567. doi:10.4161/auto.7.12.17991

41. Ferguson J, Smith M, Zudaire I, Wellbrock C, Arozarena I. Glucose availability controls ATF4-mediated MITF suppression to drive melanoma cell growth. Oncotarget. 2017;8: 32946–32959. doi:10.18632/oncotarget.16514

42. Lin T-Y, Abbassi NEH, Zakrzewski K, Chramiec-Głąbik A, Jemioła-Rzemińska M, Różycki J, et al. The Elongator subunit Elp3 is a non-canonical tRNA acetyltransferase. Nat Commun. 2019;10: 625. doi:10.1038/s41467-019-08579-2

43. Chen K, Manga P, Orlow SJ. Pink-eyed dilution protein controls the processing of tyrosinase. Mol Biol Cell. 2002;13: 1953–1964. doi:10.1091/mbc.02-02-0022

44. Toyofuku K, Wada I, Valencia JC, Kushimoto T, Ferrans VJ, Hearing VJ. Oculocutaneous albinism types 1 and 3 are ER retention diseases: mutation of tyrosinase or Tyrp1 can affect the processing of both mutant and wild-type proteins. FASEB J Off Publ Fed Am Soc Exp Biol. 2001;15: 2149–2161. doi:10.1096/fj.01-0216com

45. Fribley A, Zhang K, Kaufman RJ. Regulation of Apoptosis by the Unfolded Protein Response. Methods Mol Biol Clifton NJ. 2009;559: 191–204. doi:10.1007/978-1-60327-017-5_14

46. Manga P, Bis S, Knoll K, Perez B, Orlow SJ. The unfolded protein response in melanocytes: activation in response to chemical stressors of the endoplasmic reticulum and tyrosinase misfolding. Pigment Cell Melanoma Res. 2010;23: 627–634. doi:10.1111/j.1755-148X.2010.00718.x

47. Cheng T, Orlow SJ, Manga P. Loss of Oca2 disrupts the unfolded protein response and increases resistance to endoplasmic reticulum stress in melanocytes. Pigment Cell Melanoma Res. 2013;26: 826–834. doi:10.1111/pcmr.12158

48. Boissy RE, Liu YY, Medrano EE, Nordlund JJ. Structural aberration of the rough endoplasmic reticulum and melanosome compartmentalization in long-term cultures of melanocytes from vitiligo patients. J Invest Dermatol. 1991;97: 395–404. doi:10.1111/1523-1747.ep12480976

49. Dell’Anna ML, Ottaviani M, Albanesi V, Vidolin AP, Leone G, Ferraro C, et al. Membrane lipid alterations as a possible basis for melanocyte degeneration in vitiligo. J Invest Dermatol. 2007;127: 1226–1233. doi:10.1038/sj.jid.5700700

